# The antioxidative effect of oligomeric procyanidins on neutrophil extracellular traps alleviate chemotherapy-induced chronic kidney injury via gut-kidney axis

**DOI:** 10.1101/2024.09.03.610977

**Authors:** Yaqi Luan, Weiwei He, Kunmao Jiang, Shenghui Qiu, Lan Jin, Xinrui Mao, Ying Huang, Wentao Liu, Jingyuan Cao, Lai Jin, Rong Wang

## Abstract

Cisplatin is one of the most widely used chemotherapeutic agents for various solid tumors in the clinic, but its use is limited by adverse effects in normal tissues. In particular, cisplatin administration often damages the kidneys. However, little is known about how to alleviate cisplatin-induced chronic kidney disease (CKD) specifically. Here, we found that repeated low-dose cisplatin (RLDC) recruited susceptible neutrophils to the proximal tubule, thereby promoting the progression of CKD in the mouse model. Mechanically, cisplatin destroyed the intestinal epithelium, which induced dysregulation of gut flora and intestinal leakage. Under the dual stimulation of intestinal bacteria and cisplatin, neutrophils become extremely susceptible to form Neutrophil extracellular traps (NETs), accumulating in the proximal tubule and promoted chronic inflammation, fibrosis, and chronic hypoxia, leading to poor regeneration that promoted CKD progression. NETs provided a scaffold for tissue factors (TF) adhesion and metalloid-matrix protease 9 (MMP-9) activation, which triggered local ischemia and hypoxia. In addition, NETs promoted inflammasome construction through NOD-like receptor thermal protein domain associated protein 3 (NLRP3) shear and secretion of mature interleukin-18 (IL18), which subsequently released interferon– γ (IFN-γ), contributing to renal interstitial fibrosis. We proposed that oligomeric procyanidins (OPCs) ameliorated RLDC-induced CKD through multi-targeting damage induced by NETs. OPCs ameliorated microcirculatory disorders and inhibited inflammation by protecting the intestinal mucosa barrier and subsequent bacterial endotoxin translocation. Furthermore, we found that OPCs altered the susceptibility of neutrophils to form NETs. In summary, OPC alleviates CKD by inhibiting NETs production via TF/MMP-9 and IL-18-NLRP3 pathways. OPCs protect the kidney through anti-inflammatory and antioxidant activities, and restore the balance of the intestinal flora.

## Introduction

Cisplatin is the first platinum-based antineoplastic drug approved by the FDA (Famurewa et al., 2022). As the backbone of treatment regimens across a broad spectrum of malignancies, cisplatin improves survival and cure rates. Despite this, nephrotoxicity still limits life-saving therapy(Sahni et al., 2009). Nephrotoxicity impedes the dose and intensity of cisplatin, impacting patients’ long-term quality of life(Crona et al., 2017). To minimize side effects, clinicians often use low doses and multiple cycles of cisplatin treatment, also called repeated low-dose cisplatin (RLDC). It is more likely to induce chronic kidney disease (CKD) than acute kidney injury (AKI), as the current RLDC model shows some pathological manifestations in accordance with the critical features of CKD, including chronic inflammation(Fu et al., 2023), tubulointerstitial fibrosis(Li et al., 2023), atubular glomeruli and glomerulosclerosis(Sharp et al., 2018). However, the clear role of RLDC on CKD and the corresponding mechanism are not fully understood. In this study, we used an RLDC model (weekly repetitive low-dose cisplatin), which allowed the mice to survive 6 months posttreatment without exhibiting any clinical signs of AKI.

Neutrophils are the first leukocytes to be recruited to the sites of inflammation as the initial host defense against an extensive range of pathogens, killing harmful microorganisms in three ways: phagocytosis(Nordenfelt and Tapper, 2011), degranulation of cytotoxic enzymes(Stapels et al., 2015) and neutrophil extracellular traps (NETs), which are DNA meshes with associated cytotoxic enzymes and histones that are released into the extracellular space where they trap microorganisms(Burgener and Schroder, 2020). However, NETs are a double-edged sword; they also lead to damage, acting as a cascade to amplify and maintain inflammation. It is reported that NETs play a key role in autoimmune diseases such as lupus nephritis(Hakkim et al., 2010), diabetic kidney disease(Gupta et al., 2022), and ANCA-associated vasculitis(Yoshida et al., 2013). During NETs formation, neutrophil elastase (NE) and myeloperoxidase (MPO) facilitate nuclear membrane rupture and chromatin unwinding. Protein arginine deiminase 4 (PAD4), a histone citrullination-associated protease whose activity is greatly enhanced by binding to Ca(Lewis et al., 2015), is essential for NETs formation. Recent studies have shown that PAD4-deficient mice do not form NETs that induce thrombosis and inflammation (Leppkes et al., 2022; Raup-Konsavage et al., 2018; Seri et al., 2015). In this study, we used PAD4 knockout mice to detect the effect of NETs

NOD-like receptor thermal protein domain-associated protein 3 (NLRP3)-inflammasomes perform an essential role in inflammation. An increasing amount of data has revealed a close relationship between NETs and the NLRP3 inflammasome(Warnatsch et al., 2015; Westerterp et al., 2018). Our previous data revealed that NETs cooperated with NLRP3 to increase the levels of interleukin-18 (IL18) and interferon γ (IFNγ), which induced peripheral neuralgia after chemotherapy(Chen et al., 2020; Tapmeier et al., 2010). Alexandra and colleagues reported that cisplatin significantly induced the release of NETs with NLRP3 inflammatory vesicles involved in acute kidney injury(Mousset et al., 2023). However, NETs-NLRP3 often appear in acute inflammation, and their role in recurrent chronic inflammation is unclear. Patients with chronic kidney disease (CKD) frequently exhibit aberrant coagulation patterns(Matsushita et al., 2022). Similarly, the aggravating factors for thrombosis are positively correlated with kidney injury in a cisplatin-induced model(Watanabe et al., 2019). Aberrant coagulation can easily lead to hypoxia, which has been shown to activate hypoxia inducible factor 1α (HIF1α) to induce coagulation and increase TF levels, and HIF1α can directly inhibit TFPI promoter activity(Cui et al., 2016). Tissue factor (TF) pathway inhibitors (TFPIs) are responsible for inhibiting TF-induced coagulation. Neutrophils release elastase to cleave TFPI, counteracting its strong inhibitory effect on TF activity and subsequent thrombosis(Massberg et al., 2010). Furthermore, neutrophil-released metalloid-matrix protease 9 (MMP9)(Carmona-Rivera et al., 2015) is thought to be a HIF 1α-dependent angiogenic gene and proangiogenic protease (Song et al., 2009). Thus, we aim to investigate whether TF, TFPI, and MMP9 are involved in NETs-mediated thrombosis.

Procyanidins are known to have antibacterial, anti-inflammatory(Pallarès et al., 2013) and oxidative stress-reducing effects that can repair gut damage to reduce LPS production(Nallathambi et al., 2020). Oligomeric proanthocyanidins (OPCs) are small-molecular-weight proanthocyanidins that are generally dimers to tetramers. It has been reported that OPCs extracted from grapes improve CKD progression by reducing oxidative stress(Zhu and Du, 2020). We previously reported that OPCs strongly inhibited morphine-induced NLRP3 inflammatory vesicles in a model of gout pain(Cai et al., 2016; Liu et al., 2017). Thus, does OPCs also have an effect on NETs? Is this role vital in CKD? Our present findings showed that OPCs directly inhibited the formation of NETs, indirectly improved intestinal leakage, and ultimately alleviated RLDC-induced kidney injury.

## Results

### NETs accumulated in the kidneys after RLDC treatment

The mice were treated with 7 mg/kg cisplatin weekly for four weeks, with no significant change in mortality but a substantial decrease in body weight after 1 month of RLDC treatment (Figure 1C). Compared with those from control mice, kidneys from RLDC-treated mice presented reductions in volume and weight (Figure 1A and B), suggesting decreased repair function following renal injury after RLDC. Furthermore, pathological analysis revealed that RLDC caused dramatic changes in renal structure, including tubular dilatation and necrosis, brush border loss, tubular formation, tubular atrophy, and inflammatory cell infiltration (Figure 1D and E). The levels of serum creatinine and urea nitrogen were significantly increased (Figure 1F and G). Both plasma CitH3 and cfDNA, which are components of NETs, were elevated (Figure 1H-J), and the release of NETs were detected in the blood (Figure 1K). Numerous studies have shown the involvement of NETs in thrombosis during chemotherapy, and our previous studies demonstrated that elevated NETs in the blood of chemotherapeutic patients lead to ischemia and hypoxia in the peripheral circulation(C.-Y. Wang et al., 2023). Consistent with the above studies, Doppler flow data also revealed a significant decrease in the plantar blood flow of the mice after RLDC (Figure 1L and M). These findings suggested the increased NETs in the blood after RLDC resulted in a decrease in plantar blood flow. Furthermore, elevated CitH3 co-labeled with MPO in renal tissues suggested that cisplatin-induced NETs entered the kidneys along with the bloodstream, which was associated with kidney injury. Immunofluorescence staining revealed extensive co-localization of MPO and CITH3 in the tubular interstitium and a minor presence in the glomerular capsule. Both were associated with tubular epithelial loss (Figure 1N-Q).

**Figure 1.**
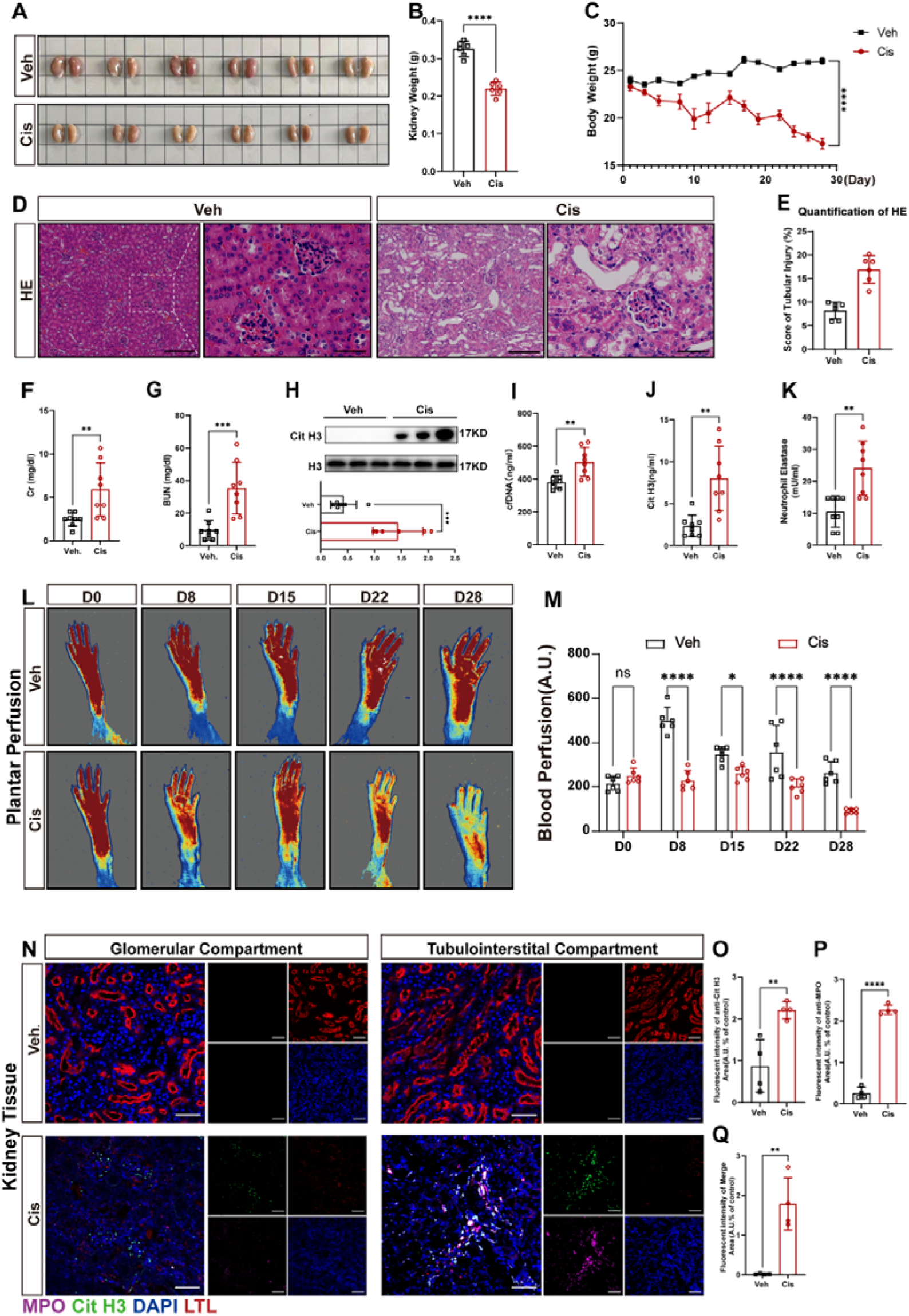
RLDC triggered NETs formation. The mice were injected weekly intraperitoneally with 7 mg/kg cisplatin for four weeks and analyzed by execution after 1 month (Cis group) or the untreated control (Veh group). Blood and kidney tissues were collected one week after the last cisplatin injection. **A**. Representative image of kidney size. **B, C.** Quantitative analysis of kidney weight and body weight (*n* = 6, **** *p* < 0.0001). **D.** Representative HE-stained images (bar = 100 μm, magn(Liu et al., 2025)ified images bar = 50 μm). **E.** Pathological tubular atrophy score (*n* = 6, **** *p* < 0.0001). **F, G.** The concentrations of serum creatinine and blood urea nitrogen (*n* = 8, \*\**p* = 0.0083, and\*\*\**p* = 0.0008). **H.** Detection of CitH3 protein levels in the kidney via western blotting (*n* = 8, \*\**p* = 0.0026). **I-K.** The contents of H3Cit, NE, and cfDNA in plasma after intraperitoneal injection of cisplatin were evaluated via the H3Cit ELISA kit, NETosis assay, and dsDNA ELISA kit (*n* = 8, \*\**p* < 0.01). **L, M.** Blood flow in the lower limbs of the mice was measured at the start of cisplatin treatment and at the end of each week via a MoorFLPI2 blood flow scatter hemodynamometer. (*n* = 6, \*\*\*\**p* < 0.0001, D28). **N-Q.** NETs (confocal immunofluorescence microscopy images; stained for MPO, Cit H3, LTL, and DNA) visible in kidney samples from chemotherapeutic nephritic mice (*n* = 4, \*\**p* < 0.01, \*\*\**p* < 0.001). Scale bar, 50 μm. Significant differences were revealed via one-way ANOVA *vs.* vehicle (A, B, E, F-K, M, and O-Q).

### Blocking NETs prevented RLDC-induced renal tubular injury and apoptosis

We proceeded to investigate whether the presence of NETs was involved in kidney injury. It is an essential process for PAD4 to enter neutrophils, where PAD4 citrullinates histone arginine residues to deconcentrate chromatin, generating NETs. Thus, PAD4 knockout (PAD4-/-) mice, which are not able to form NETs, were used. As shown in Figure 2A and B, RLDC induced renal atrophy and body weight loss in both WT and PAD4−/− mice. However, in contrast to the WT mice, of which kidneys presented marked brush border loss, cast formation, massive loss of tubular epithelial cells, tubular dilatation, and tubular intratubular debris, PAD4 knockout inhibited tubular injury, with well-preserved brush border membranes and no loss of tubular epithelial cells (Figure 2C). The collagen accumulation in the renal interstitial and perivascular areas of WT mice were obvious, while PAD4 knockout significantly reduced it. Moreover, the absence of PAD4 attenuated interstitial fibrosis of the kidneys (Figure 2D and E). Further semiquantitative analysis of HE, PAS, and Masson’s trichrome staining revealed that tubular injury, collagen deposition, and tubulointerstitial fibrosis scores were significantly higher in RLDC-treated WT mice, whereas the scores in RLDC-treated PAD−/− mice were similar to those in untreated mice (Figure 2G-I). Under RLDC treatment, WT mice presented proximal renal tubular degeneration with loss of the brush border and LTL tubules, accompanied by increased KIM-1, which disappeared in PAD4−/− mice (Figure 2F). As shown in the semiquantitative analysis, the percentage of the KIM-1-positive area decreased from 2·43% to 0·58%, and the LTL-positive area recovered from 2·39% to 6·68% after PAD4 deficiency (Figure 2J and K). Consistent with the improvement in renal function, the expression of KIM-1 and BAX was increased in WT mice but not in PAD4−/− mice, suggesting that PAD4 was involved in CKD induced by RLDC (Figure 2L-N).

**Figure 2.**
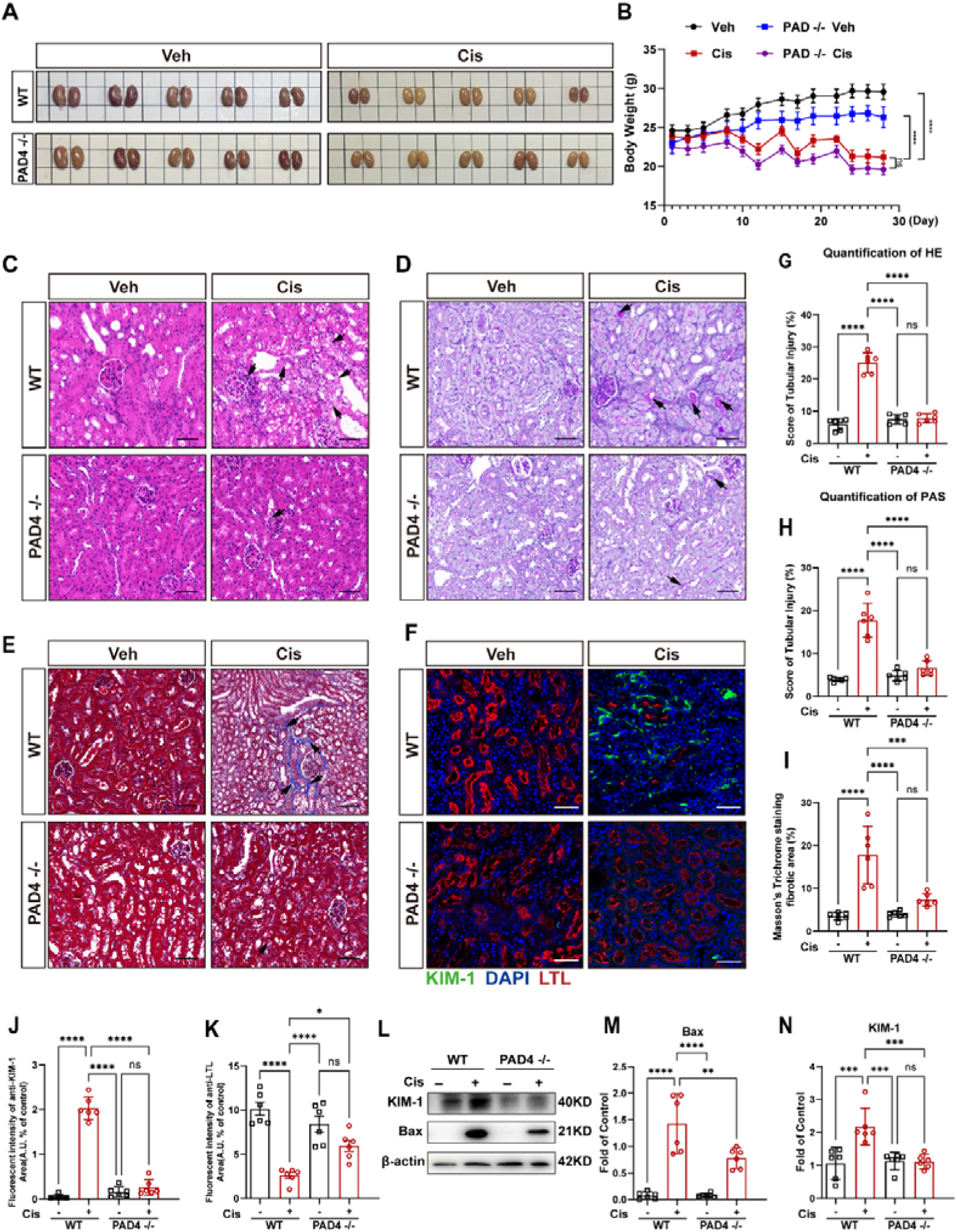
NETs mediated kidney injury. **A.** Gross observation of kidneys from RLDC-induced WT or PAD4−/− mice. A representative image of kidney size is shown (*n* = 5). **B.** Trends in the body weights of the mice during the four weeks of modeling (*n* = 10). **C-E.** HE, PAS, and Masson’s trichrome staining of kidney slices (scale bars 50 μm, *n* = 6). **F.** Representative confocal images of KIM-1+ and LTL+ tubules (scale bars 50 μm, *n* = 6). **G-I.** Quantification of renal tubular damage and renal fibrosis in the mice in each group (n = 6, \*\*\*\**p* < 0.0001). **J, K.** Quantification of KIM-1-positive and LTL-positive areas of the kidney (*n* = 6, \*\*\*\**p* < 0.0001). **L**O**N.** WB analysis and quantification of KIM-1 and BAX expression in the kidney in each group as indicated (*n* = 6, \*\*\**p* < 0.0001, and \*\*\*\**p* < 0.0001). Significant differences were revealed via one-way ANOVA *vs*. Vehicle/PAD4−/− (B, J-K, M, and N).

### Blocking NETs prevented the activation of NLRP3 inflammatory vesicles

Our previous study demonstrated that NETs induced the maturation of NLRP3 inflammatory vesicles(Lin et al., 2022). To investigate whether the NLRP3 inflammasome was involved in NETs-mediated renal injury, the levels of NLRP3 and related inflammatory factors were detected in PAD4−/− mice. Firstly, with RLDC treatment, the mice exhibited severe renal dysfunction, as evidenced by elevated plasma creatinine and urea nitrogen, which were significantly lower in the PAD4−/− mice (Figure 3A and B). The levels of NETs are very low in both serum and kidneys of PAD4−/− mice (Figure 3C-F, L). Secondly, western blot analysis revealed that RLDC increased the expression of NLRP3, Casp-1, interleukin 1β (IL1β), interleukin 18 (IL18), and interferon γ (IFNγ) (Figure 3F-K). IL18, a Th1 cytokine, significantly induces natural killer (NK) cells and T cells to produce interferon gamma (IFNγ), which is involved in cytotoxicity and type I immunity(Landy et al., 2024). In addition to its well-known antifibrotic effects, IFNγ is involved in the repair of renal tubulointerstitial fibrosis. This is a double-edged sword, as it can also induce fibrosis(Kim et al., 2022). As demonstrated by immunofluorescence, IFNγ was present in almost every lumen of LTL-labeled tubules following RLDC (Figure 3M). Compared with wild-type mice, the positivity rate of IFNγ in PAD4−/− mice is only 20.54% of that in wild-type mice (Figure 3O). As expected, RLDC induced renal tubulointerstitial fibrosis in WT mice, accompanied by an increase in α-SMA. These changes were not obvious in PAD4−/− mice (Figure 3N and P). However, following the administration of NLRP3 and IL-18 inhibitors, the survival rate did not show significant improvement under the stimulation of RLDC. Furthermore, the function of the mice’s kidneys was worse than that of the model group in the third week of the modeling (Supplementary Figure 3). This finding indicates that the major pathogenic effect of NETs on CKD in this study may not be the NLRP3-IL18 pathway.

**Figure 3.**
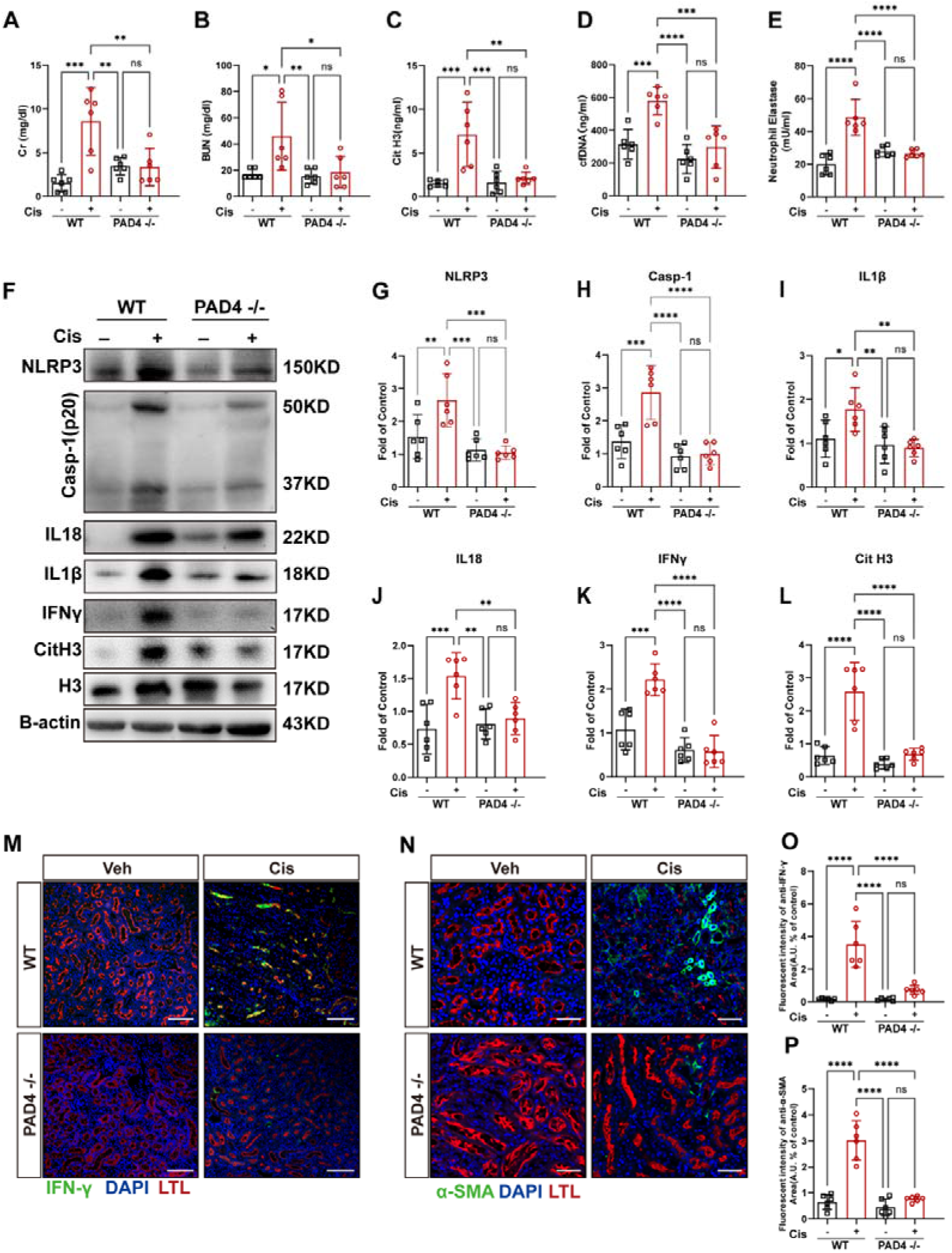
NETs activate the NLRP3 inflammasome and subsequent renal fibrosis. **A, B.** Induction of CKD in WT and Pad4−/− mice treated with 7 mg/kg cisplatin for four weeks. Serum creatinine and blood urea nitrogen (*n* = 6, \*\*\**p* < 0.0001, and \*\**p* = 0.0049). **C-E.** The contents of Cit H3, NE, and cfDNA in plasma after intraperitoneal injection of cisplatin were evaluated via the Cit H3 ELISA kit, NETosis assay, and dsDNA ELISA kit (*n* = 6, \*\*\*\**p* < 0.0001). **F**O**L.** Western blot analysis of NLRP3, Casp-1, IL18, IL-1β, IFNγ and Cit H3 in kidney tissues. β-actin was used as a loading control (*n* = 6, \*\**p* = 0.0029, \*\*\**p* = 0.0004, \*\*\*\**p* < 0.0001). **M.** Representative images of costaining for IFNγ (green), LTL (red), and DAPI (blue). Scale bar, 100 μm. **N.** Representative images of costaining for α-SMA (green), LTL (red), and DAPI (blue). Scale bar, 50 μm. **O, P.** Quantification of IFNγ-positive and α-SMA-positive areas in the kidney (*n* = 6, \*\*\*\**p* < 0.0001). Significant differences were revealed via one-way ANOVA *vs.* vehicle/PAD4−/−(A-E, J-L, O and P).

### NETs-mediated functional TF-MMP9 activation was required for renal ischemia and hypoxia

Hypoxia plays a key role in the pathogenesis of CKD and promotion of coagulation and thrombotic events(Oe and Takahashi, 2022). A recent study demonstrated that hypoxia-inducible factor 1 alpha (HIF-1α) was involved in renal fibrosis in the RLDC mouse model(Zhao et al., 2021). We found that RLDC led to microcirculation disorders in this study (Figure 1L). To explore whether RLDC-induced NETs cause microthrombi in the kidneys and their roles in renal tubular injury, we examined the levels of HIF1α and relevant microthrombus indicators. As shown in Figure 4A-E, RLDC significantly elevated the levels of HIF-1α, TF, and MMP9 and reduced the levels of TFPI in the renal tissues of WT mice. In contrast, RLDC had no discernible effect on PAD4−/− mice. Surprisingly, most NETs colocalized with TF in the renal tubular mesenchyme at the corticomedullary junction, which was also the most sensitive to hypoxia, where there were more marked pathological changes in a state of hypoxia. In contrast, due to the absence of NETs, no obvious TF was deposited in the PAD4−/− mice (Figure 4F-H). Long-term hypoxia usually results in interstitial hyperplasia. Ki67 is a marker of cell proliferation reflecting hyperplasia. A significant increase in the number of Ki67-positive cells was observed in the kidneys of WT mice following RLDC treatment, with a particularly notable increase in the region of the junction between the renal cortex and the medulla (Figure 4I). In contrast, the number of Ki67-positive cells in PAD4−/− mice decreased by 55·29% (Figure 4K), suggesting that NETs cooperating with TF stimulated interstitial hyperplasia. Finally, we employed laser Doppler speckle flowmetry to assess plantar blood flow in the mice. Compared with that in WT mice, the mean lower limb blood flow in PAD4−/− mice was not influenced by RLDC (Figure 4J and L). This finding aligns with the hypothesis that NETs contribute to the exacerbation of thrombus formation.

**Figure 4.**
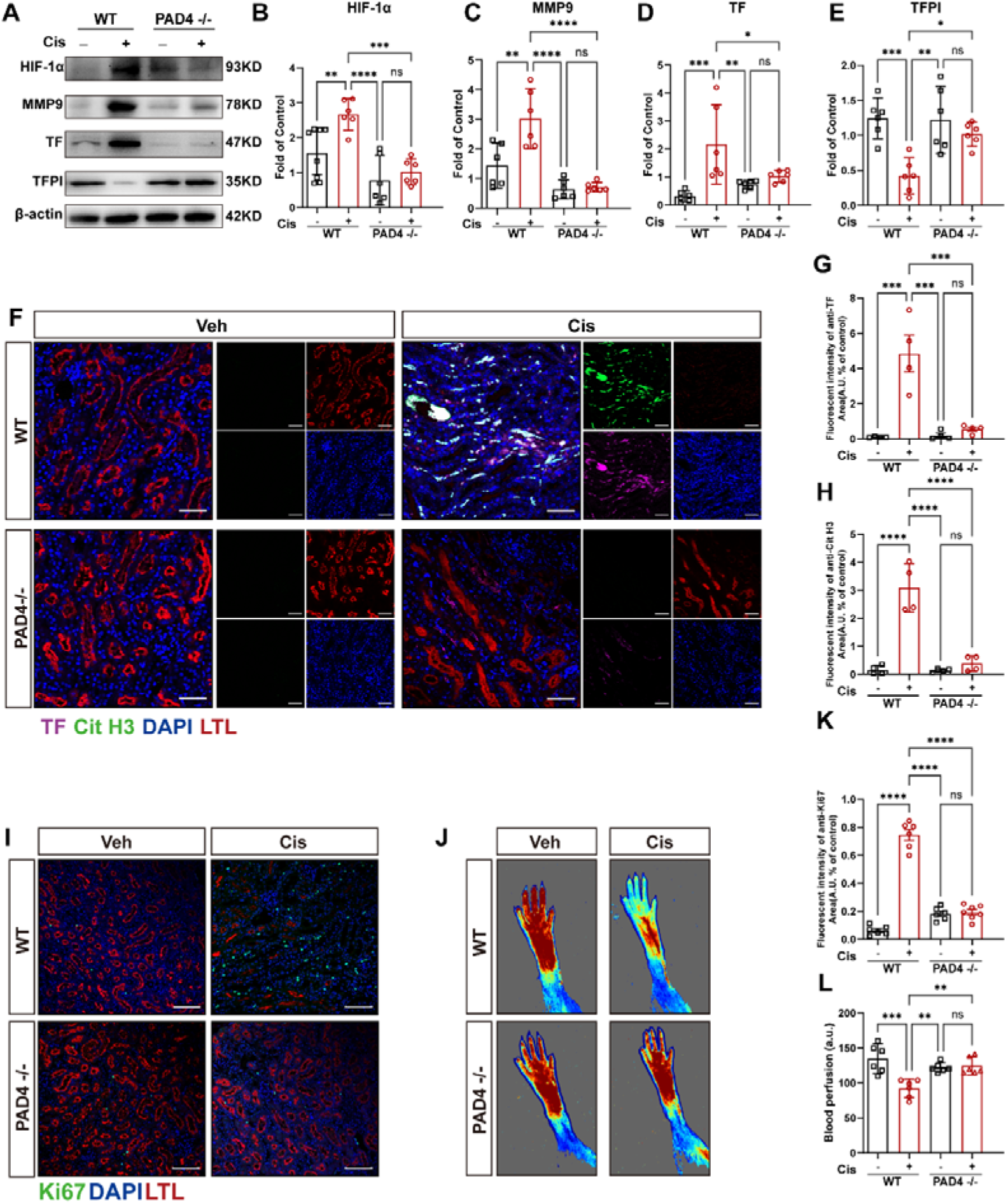
NETs trigger thrombus formation, leading to local ischemia and hypoxia. **A-E**. Western blot for HIF-1α, TF, MMP9 and TFPI in kidney tissues. β-actin was used as a loading control (*n* = 6, \*\**p*=0.0012, and \*\*\**p*=0.0004, \*\*\*\**p* < 0.0001). **F.** Confocal immunofluorescence microscopy images of kidney samples from chemotherapeutic nephritic mice (*n* = 4) stained for TF, Cit H3, LTL and DNA. Scale bar, 50 μm. **G, H.** Quantification of Cit H3-positive and TF-positive areas in the kidney (*n* = 4, \*\*\*\**p* < 0.0001). **I.** Representative images of Ki67 immunofluorescence staining and costaining with LTL. Scale bar, 100 μm. **K.** Quantification of the Ki67 area in the kidney (*n* = 6, \*\*\*\**p* < 0·0001). **J, L.** Measurement of lower limb blood flow in WT and PAD4−/− mice via a MoorFLPI2 blood flow scattering hemodynamometer (*n* = 6, \*\*\**p* = 0.0003). Significant differences were revealed via one-way ANOVA *vs.* vehicle/PAD4−/−(B-E, G, H, K and L).

### OPCs inhibited LPS leakage caused by intestinal barrier damage

Cisplatin causes severe damage to the intestinal mucosa, disrupting the intestinal mucosal barrier, leading to leaky gut, and causing long-term chemotherapeutic colitis(Hu et al., 2021). These findings prompted us to speculate that intestinal damage might contribute to cisplatin-induced CKD. To investigate whether OPCs can alleviate cisplatin-induced CKD through the protection of the intestine, we treated the mice with OPCs and used berberine (BBR) as an anti-inflammatory and antibacterial positive control.(Chen et al., 2017; Pallarès et al., 2013; Pan et al., 2023; Zhang et al., 2019; Zhu et al., 2022) As shown in Figure 5A and B, the intestinal length was significantly decreased after RLDC treatment, but was restored by BBR or OPCs. RLDC resulted in the disappearance of villi due to atrophy and detachment, as well as inflammatory cell infiltration, in addition to the disappearance of glands, cup cells and crypts. Notably, inflammatory cell infiltration was absent in both the BBR and OPCs treatment groups. OPCs effectively restored the intestinal villi (Figure 5C and D), suggesting that both BBR and OPCs significantly inhibited cisplatin-induced colonic inflammation. We also used laser Doppler imaging to detect the intestinal blood flow of mice. RLDC significantly reduced intestinal blood flow, which was restored by both BBR and OPCs (Figure 5E and F). To further assess the status of intestinal damage, the protein expression of the intestinal barrier was quantitatively analyzed. The expression of Claudin-1, ZO-1, and Occludin were reduced by RLDC but reversed by BBR or OPCs (Figure 5G-L, Figure 5J-M). Next, we detected plasma LPS in the mice. Elevated LPS after RLDC suggested that intestinal leakage and bacteria entered the bloodstream. BBR and OPCs attenuated the level of cisplatin-induced LPS (Figure 5N). In addition, IVIS spectrum imaging directly revealed that RLDC stimulated FITC-dextran leakage from the intestine and elevated serum FITC-dextran levels, which were prevented by both OPCs and BBR (Figure 5O and P). These results indicate that OPCs might play a role in preventing cisplatin-associated CKD by restoring the intestinal barrier.

**Figure 5.**
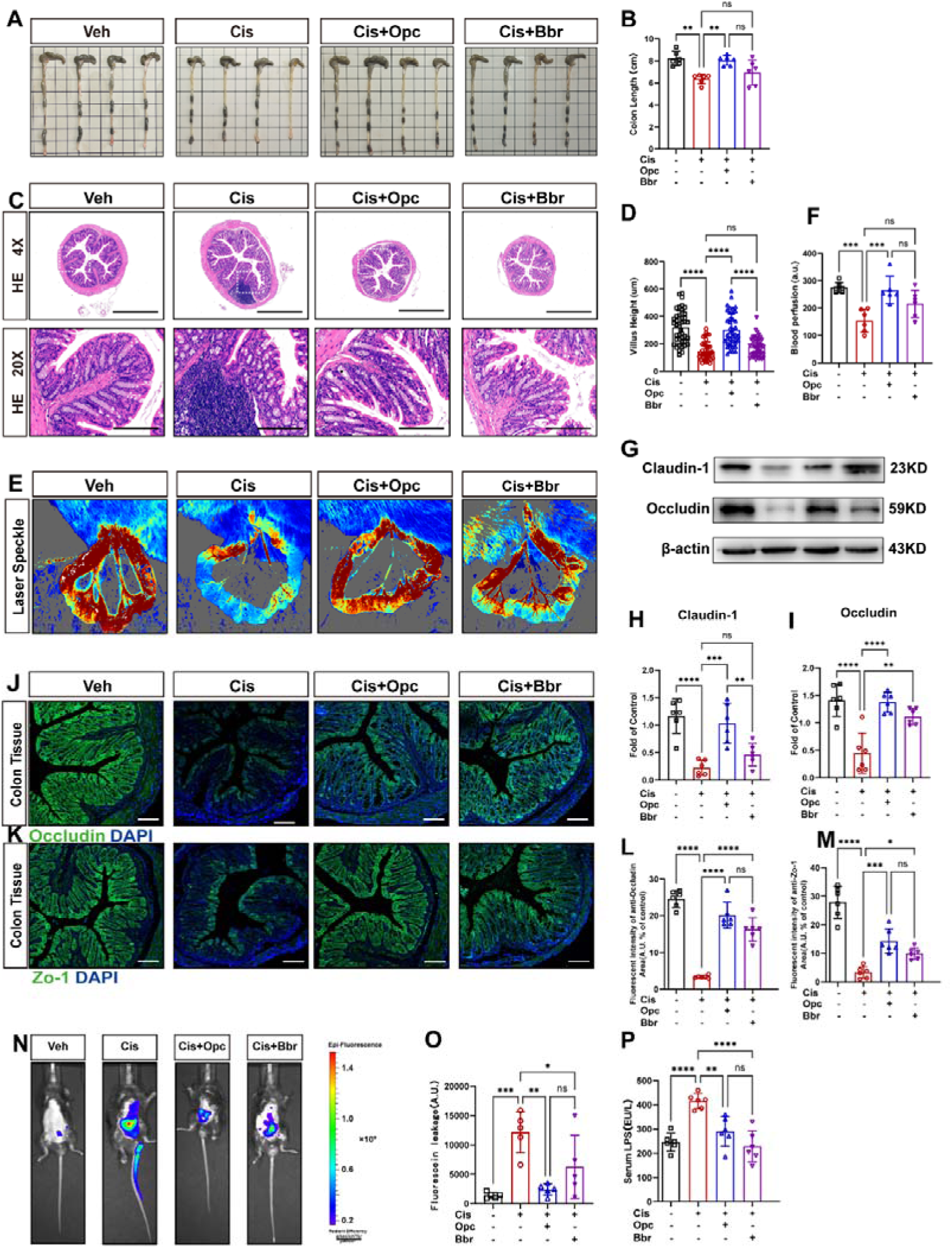
OPCs retain the integrity of the cisplatin-treated intestinal barrier. **A, B.** Macroscopic images and the length of the colon from each group were measured (*n* = 6, \*\*\**p* = 0·0005). **C.** HE staining of colon sections (scale bars 50 μm, *n* = 6). **d.** Colonic villus length (40 villi per group, \*\*\*\**p* < 0.0001). **E, F.** Intestinal blood flow and perfusion indices were measured via a laser speckle blood flow analysis system (*n* = 6). **G-I.** Immunoblot analysis of tight junctions in the colons of cisplatin-treated mice (*n* = 6, \*\*\*\**p* < 0.0001). **J-M.** The expression levels of the tight junction proteins ZO-1 and Occludin were observed via immunofluorescence (scale bars 100 μm, n = 6, \*\*\*\**p* < 0·0001). **N.** FITC-dextran distribution in the mice with colitis was observed via small animal imaging. **O.**Content of FITC-dextran in serum (*n* = 5, \*\*\**p* = 0.0003). **P.** LPS levels in the serum of the mice (*n* = 6, *p* < 0.0001).Significant differences were revealed via one-way ANOVA (B, D, F, H, I, K, M, N, and P).

### OPCs maintained intestinal flora homeostasis

The destruction of the intestinal barrier by cisplatin may be accompanied by disruption of the gut microbiota. We investigated whether cisplatin altered the composition of the gut flora via 16S rDNA gene sequencing. Firstly, there were significant changes in the Chao1 index, Shannon index, phylogenetic diversity (PD), and species of bacteria in the cisplatin group, demonstrating that RLDC significantly reduced the species diversity and homogeneity of the intestinal flora, whereas OPCs restored the richness of the intestinal flora (Figure 6A-D). Moreover, intergroup analysis by Metastats revealed a significant difference between the control and cisplatin groups, indicating that RLDC led to a clear separation between the biological communities (Figure 6E). β diversity analysis was subsequently performed, and unweighted UniFrac distance PCoA revealed that the composition of the intestinal flora was completely different between the control and cisplatin groups (Figure 6F). The composition of the OPCs-treated group was intermediate, suggesting the unique ability of OPCs to regulate the intestinal flora, which may be related to the mechanism of OPCs in CKD. To further confirm the specific composition of the intestinal flora in the RLDC state, we analyzed the intestinal flora at each group’s phylum and genus levels. The abundances of Firmicutes and Bacteroidetes were lower in the cisplatin group. In contrast, OPCs group contained an abundance of Bacteroidetes. The Firmicutes/Bacteroidetes (F/B) ratio is reportedly related to the microbiological state of the gut in chronic colitis(Kieffer et al., 2016). Unfortunately, OPCs did not obviously affect this ratio (Figure 6G and H). Based on absolute species abundance information, Muribaculaceae, Akkermansia, and Clostridia-UCG-014 declined after RLDC treatment and were significantly recovered by OPCs (Figure 6L). The same evolutionary map generated via LEfSe revealed differences among the three groups of taxa (from phylum to genus), with the model significantly different from the control. The dominant group present in the OPCs was c_Clostridia p_Firmicutes (Figure 6J and K). We also performed TAX4fun analysis to predict gut microbial function. CIS was enriched in several pathways, including the nervous system metabolism of cofactors and vitamins, whereas it inhibited several functions, including the cellular community, circulatory system, and signaling molecules. OPCs restored, as much as possible, the original function of the gut flora (Figure 6I). According to FAPROTAX functional analysis, xylanolysis and sulfate respiration were decreased, and hydrocarbon degradation and aliphatic non-methane hydrocarbon degradation were elevated in the Mod group, whereas these changes were reverted to the usual by OPCs (Figure 6M). To investigate the effects of OPCs on potential metabolic pathways in the intestinal microbiota, a t-test was conducted on the annotated abundance data of secondary pathways within the KEGG metabolic pathway for a specific set of two samples. As shown in Figure 6N, the metabolism of the intestinal flora in the model group was markedly aberrant, e.g., reduced carbohydrate metabolism and increased amino acid metabolism. In conclusion, the significant changes mentioned above confirmed the modulatory effect of OPCs on intestinal flora homeostasis, which resisted the intestinal damage induced by RLDC.

**Figure 6.**
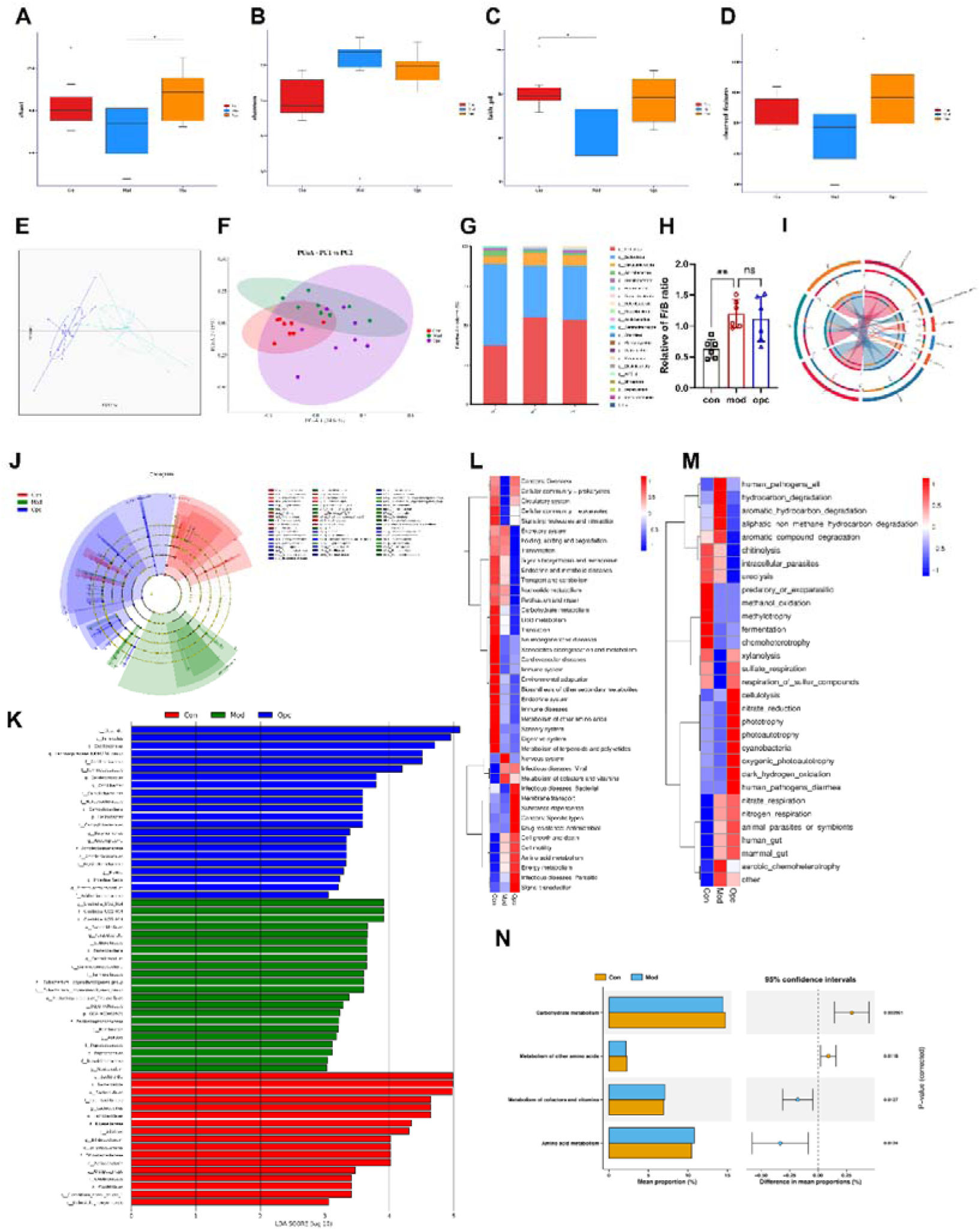
Analysis of the mouse gut microbiota via 16S rDNA gene sequencing. **A-D.** α-Diversity indicated by the Chao index, Shannon index, PD_whole_tree index and observed features (interquartile range, IQR, *n* = 8, \**p* < 0.05). **E.** Principal component analysis (PCA) revealed significant differences at the phylum level. **F.** Effect of OPCs on the β diversity of the gut microbiota assessed via principal coordinate analysis (PCoA). **G, H.** Relative abundance of Bacteroidetes and Firmicutes at the phylum level (*n* = 6, \*\*\**p* = 0.0002). **I.** Abundances of the top 5 species at different taxonomic levels and absolute species abundance information on the basis of ASVs within samples. **J.** LDA analysis. **K.** LEfSe analysis. **L, M.** Classification of functions based on Tax4Fun and FAPROTAX analysis. **N.** Analysis of differences in KEGG metabolic pathways. (*n* = 8, \**p* < 0.05).

### OPCs reduced NETs production by maintaining neutrophil homeostasis

In addition to restoring intestinal flora stability, the part of OPCs that is absorbed into the blood may contribute to the stability of neutrophils. Following RLDC treatment, there was an increase in the number of peripheral blood neutrophils, which was reduced by OPCs, though not to a significant difference (Supplementary Figure 2). It may be attributed to the function of neutrophils as acute response cells, being utilized in the context of chronic diseases(Liew and Kubes, 2019). As demonstrated in Supplementary Figure 1A, we extracted neutrophils from the peripheral blood of mice treated with or without OPCs from the RLDC treatment group. Peripheral blood neutrophils from the RLDC group of mice exhibited a higher level of NETs following one hour of culture. Meanwhile, the level of NETs in the OPC group was relatively low (Supplementary Figure 1B, I-M). In line with this observation, one hour after extraction, the peripheral blood from RLDC mice exhibited elevated levels of MPO and Cit H3. However, there is no significant increase in the OPCs co-administered group (Supplementary Figure 1C-F). A significant increase of ROS in the neutrophils was present in the peripheral blood of mice from the RLDC group (Supplementary Figure 1G and H). It suggests the potential involvement of ROS in the production of neutrophil NETs in the experimental model employed, and OPCs may inhibit NETs production by means of suppressing ROS. Furthermore, the re-administration of low-dose cisplatin and/or OPCs to the extracted peripheral blood neutrophils showed that OPCs enhanced the resistance of neutrophils to low-dose cisplatin (Supplementary Figure 5). To determine whether LPS from the gut was also a factor contributing to the formation of NETs, neutrophils were isolated from the bone marrow of mice with no other treatment. Interestingly, neither LPS nor cisplatin alone induced NETs formation, while co-administration of both resulted in many NETs (Figure 9A). OPCs significantly reduced the number of NETs at the same stimulus intensity (Figure 9B), further demonstrating the direct inhibitory effect of OPCs on NETs formation.

### OPCs prevented RLDC-induced CKD by inhibiting the formation of NETs

To investigate whether OPCs inhibited cisplatin-induced CKD through NETs, kidneys were collected from RLDC mice treated concomitantly with OPCs (100 mg/kg). The administration of OPCs alone did not influence any renal functions, suggesting that OPCs did not exacerbate renal stress (Supplementary Figure 4). As expected, OPCs significantly decreased the expression of NETs marker Cit-H3 (Figure 7A and M); the ischemia– and hypoxia-related factors HIF-1α, MMP9, and TF (Figure 7A-D); and the inflammation– and fibrosis-related factors NLRP3, Casp-1, IL1β, IL18, and IFNγ (Figure 7A, G, and I-K), all of which were increased by RLDC. In addition, the expression of TFPI, which was inhibited by RLDC, was elevated by OPCs. OPCs also inhibited renal injury factors, such as caspase-1, BAX and KIM-1, suggesting alleviation of ischemiaLJhypoxia and inflammationLJfibrosis (Figure 7ALJL). OPCs reduced the deposition of serum NETs (Figure 7M-P). Consistent with this trend, there was a notable decline in the serum creatinine and urea nitrogen levels. (Figure 7Q and R). Furthermore, renal tubular atrophy was significantly reduced by OPCs. The collagen area and degree of fibrosis in senescent kidneys were significantly decreased concurrently (Figure 7S-X).

**Figure 7.**
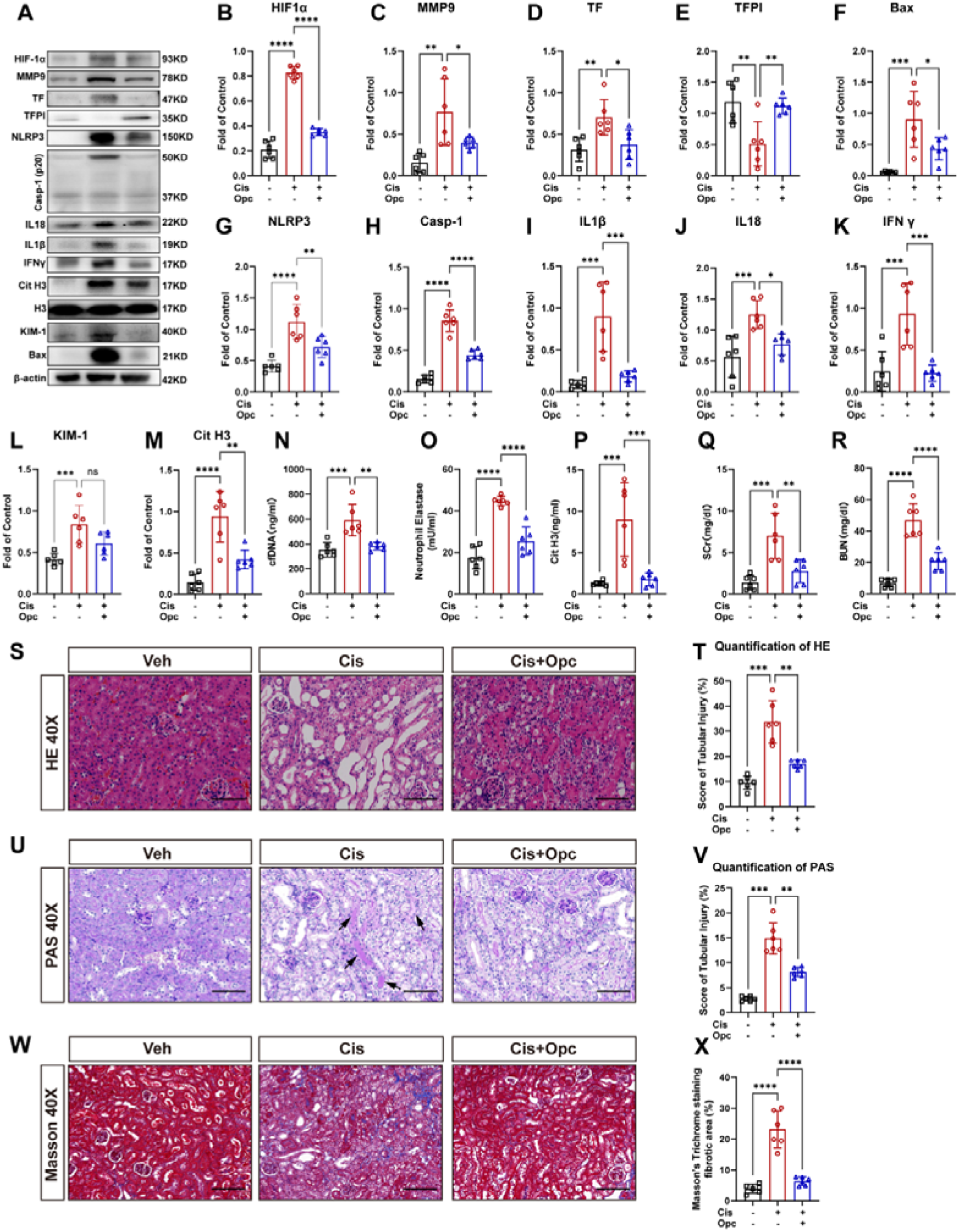
OPCs reduces kidney damage by inhibiting NETs. The administration of OPCs (100 mg/kg, i.g./3 d) was conducted in conjunction with the initial intraperitoneal cisplatin injection. RLDC treatment was administered with or without OPCs treatment and was administered one week after the last cisplatin treatment. **A**O**M.** Immunoblot analysis of HIF-1α, TF, MMP9, TFPI, BAX, NLRP3, Casp-1, IL18, IL1β, IFNγ, Cit H3 and KIM-1 in kidney tissues. For quantification, the protein was analyzed through densitometry and then normalized to β-actin (*n* = 6, \*\**p* < 0.01, \*\*\**p* < 0.001, \*\*\*\**p* < 0.0001). **N-P.** The content of Cit H3, NE, and cfDNA in plasma was evaluated via the Cit H3 ELISA kit, NETosis Assay, and dsDNA ELISA kit (*n* = 6, \*\*\**p* < 0.001, \*\*\*\**p* < 0.0001). **Q, R.** Serum creatinine (SCr) and blood urea nitrogen (BUN) levels (*n* = 6, \*\*\**p* = 0.0002, \*\*\*\**p* < 0.0001). **S.** Representative HE-stained kidney slices (scale bars = 50 μm). **T.** Quantification of renal tubular damage in the mice in each group (*n* = 6, \*\*\*\**p* < 0.0001). **U.** Representative images of PAS-stained kidney slices (scale bars = 50 μm). **V.** Quantification of renal tubular damage in the mice in each group (*n* = 6, \*\*\*\**p* < 0.0001). **w.** Masson’s trichrome staining of kidney cortex sections (scale bars = 50 μm). **X.** Quantification of the collagen-positive area according to Masson staining (*n* = 6, \*\*\*\**p* < 0.0001). Significant differences were revealed via one-way ANOVA *vs.* Vehicle/Cis+OPCs (B-R, T, V, and X).

RLDC elevated renal KIM-1 expression, which was restored by OPCs. However, LTL expression was not significantly restored by OPCs (Figure 8M-N). This may be attributed to the loss of the proximal tubular brush border as a consequence of long-term renal injury(Kishi et al., 2019; Livingston et al., 2023). OPCs also inhibited α-SMA deposition (Figure 8P and Q), thereby alleviating the susceptibility to fibrosis produced by repeated kidney injury, which is a significant factor in the development of CKD in older individuals(Ferenbach and Bonventre, 2015). OPCs inhibited the deposition of NETs (Figure 8A, Figure 8C-E) and TF-NETs microthrombosis (Figure 8B, Figure 8F-H) in the kidney induced by RLDC. Since the effect of OPCs on the kidney was better than that of berberine (Figure 5), we speculated that OPCs not only works through the intestine but also has antioxidant effects on neutrophils directly. As shown in Figure 8I and J, OPCs suppressed the cisplatin-increased ROS in neutrophils. The in vivo experiments demonstrated that OPCs restored the activities of SOD and GSH enzymes and alleviated the oxidative stress caused by long-range cisplatin (Figure 8K and L). The findings above suggested that OPCs protected kidney via anti-NETs, anti-ischemia hypoxia, anti-inflammation and anti-oxidantion.

**Figure 8.**
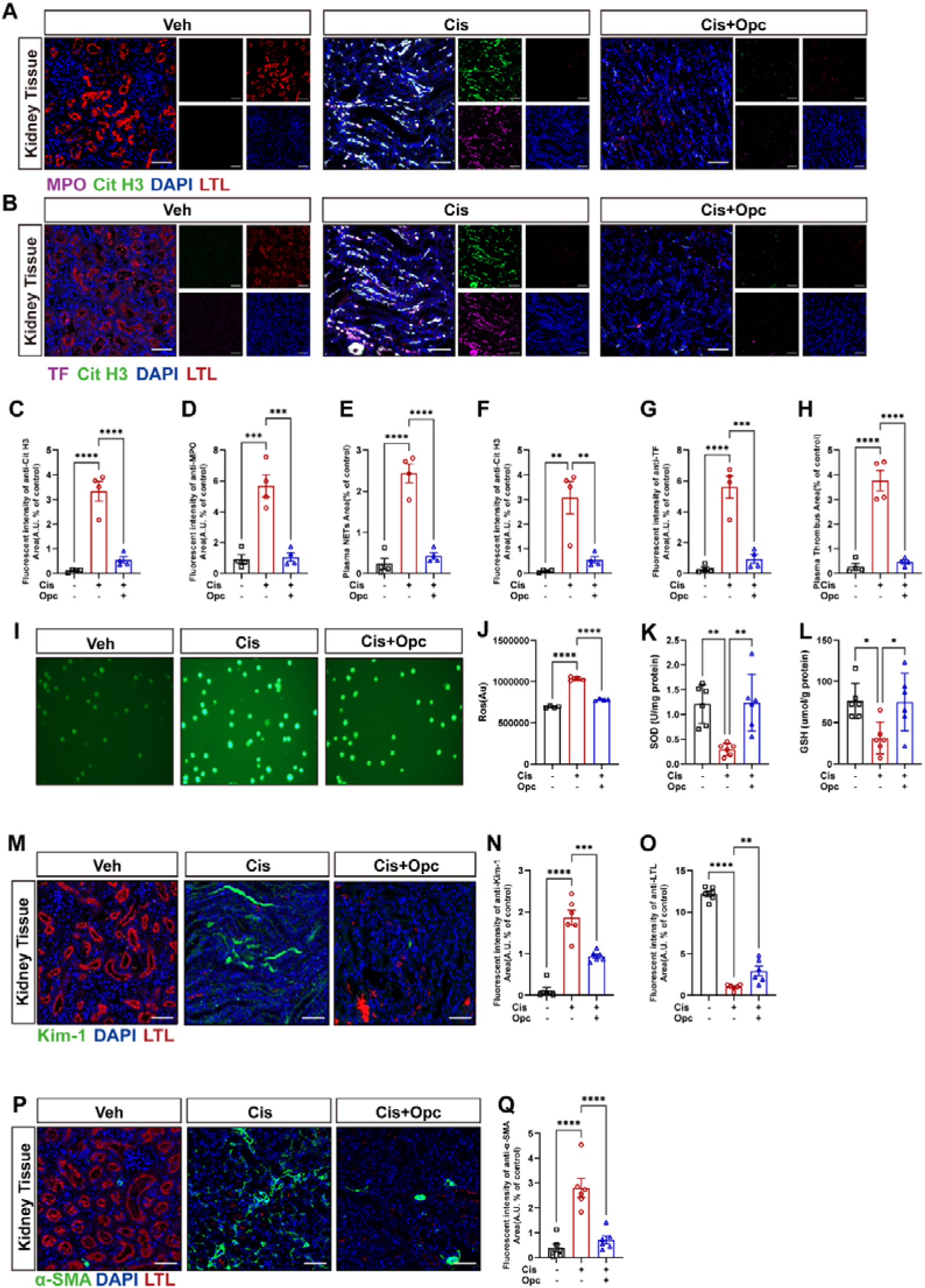
OPCs inhibit NETs production, with subsequent thromboembolism and inflammatory fibrosis. **A.** NETs (confocal immunofluorescence microscopy images; stained for MPO, Cit H3, LTL and DNA) visible in kidney samples from chemotherapeutic nephritic mice (scale bar = 50 μm). **B.** Thrombus (confocal immunofluorescence microscopy images; stained for TF, Cit H3, LTL and DNA) visible in kidney samples from chemotherapeutic nephritic mice (scale bar = 50 μm). **C-E.** Quantification of renal Cit H3 positivity, MPO positivity and the colabeled area of the kidney (*n* = 4, \*\*\*\**p* < 0.0001). **F-H.** Quantification of renal Cit H3 positivity, TF positivity and colabeled area of the kidney (*n* = 4, \*\*\*\**p* < 0.0001). **I.** ROS analysis by DCFH-DA staining in neutrophils pretreated with OPCs for 1 h followed by cisplatin for 12 h. **J.** Quantification of DCFH-DA staining of neutrophils by luminescence zymography (*n* = 4, \*\*\*\**p* < 0.0001). **K.** Serum SOD levels (n = 6, \*\**p* = 0.0015). **L.** Serum GSH levels (n = 6, \**p* = 0.0142). **m.** Representative images of KIM-1– and LTL–stained sections of mouse kidneys (scale bar = 50 μm). **n, o.** KIM-1-positive and LTL-positive areas of the kidney (*n* = 6, \*\*\*\**p* < 0.0001, \*\**p* = 0.0077). **P.** Representative images of costaining for α-SMA (green), LTL (red), and DAPI (blue). Scale bar = 50 μm. **Q.** Quantification of the α-SMA-positive area of the kidney (*n* = 6, \*\*\*\**p* < 0.0001). Significant differences were revealed via one-way ANOVA *vs.* Vehicle/Cis+OPCs (C-H, J-L, N, O, and Q).

In order to assess the toxicity of cisplatin-induced NETs on renal tubular cells, as demonstrated in Supplementary Figure 6A, tubular epithelial cells (TECs) were co-cultured with NETs for a period of 24 hours. Firstly, in comparison with low-dose cisplatin, NETs appeared to be more capable of causing TEC damage and increasing KIM-1 expression (Supplementary Figure 6B and C). Secondly, unstimulated neutrophils from the bone marrow cavity were treated with cisplatin alone or along with OPCs, the NOX inhibitor NAC or the potent NETs scavenger DNase I (Supplementary Figure 7A). Immunofluorescence revealed that NETs adhered to TEC and were engulfed by TEC, leading to increased KIM-1 expression, suggesting that NETs generated by cisplatin damaged TEC. By contrast, the KIM-1 expression in TEC treated with OPC or DNase I was significantly reduced (Supplementary Figure 7F-H), accompanied with a significant reduction in BAX expression (Supplementary Figure 7B and C).Concurrently, DCFH-DA staining revealed that ROS level was increased following cisplatin treatment, while ROS level was decreased after treatment with OPCs, NAC, or DNase I (Supplementary Figure 7D and E).. Taken together, these results suggest that inhibition of NETs formation via ROS attenuates RLDC-induced kidney damage.

## Discussion

Patients who receive multiple cycles of cisplatin chemotherapy may suffer from CKD(Latcha et al., 2016). However, the underlying mechanisms and corresponding drugs for alleviating CKD are largely unknown. In this study, we employed PAD4−/− mice to demonstrate the involvement of NETs in RLDC-induced CKD. Mechanistically, Cisplatin disrupted the intestinal barrier, resulting in the leakage of intestinal flora and subsequent sustained LPS-induced endotoxemia, which led to continuous induction of NETs. Furthermore, we elucidated the roles of the NETs-TF-MMP9 and NETs-NLRP3-IFNγ pathways in CKD. In the context of cisplatin stimulation, OPCs had the antioxidative capacity to improve renal dysfunction and tubular injury, accompanied by decreased micro-thromboembolism and chronic inflammation, consistent with the data of NETs inhibition. Notably, NLRP3 inhibitor didn’t show any significant effect in our study, suggesting elevated levels of NETs in the kidney result in TF-MMP9-associated inflammation on the one hand and ischemia and hypoxia on the other, which ultimately lead to apoptosis and interstitial fibrosis in renal parenchymal cells.

It is reported that neutrophils entering the bloodstream typically dissipate within 12 hours, exhibiting a short lifespan and extremely low concentrations in the kidneys(D et al., 2024; M et al., 2018). Consequently, resident neutrophils in the kidneys do not constitute the primary cause of local injury in CKD progression. Neutrophils represent the predominant cells in blood, constituting the primary host defense within blood vessels(Bg et al., 2017). Within the kidneys, a response to inflammatory stimuli is initiated, whereby they reside in the glomerular capillary network to form inflammatory thrombus formation(M et al., 2019). In the context of physiological conditions, the migration of neutrophils is impeded by endothelial cell gap junctions and sialic acid residues. However, it is important to note that prolonged chemotherapy has been shown to disrupt this structure, leading to significant neutrophil infiltration into the interstitium. In the context of ischemia-reperfusion-induced acute kidney injury (IRI-AKI), renal tubular epithelial cells initially respond to hypoxia via a RIPK-mediated process of apoptosis, which results in the release of HMGB1. This has been shown to trigger the release of damage-associated molecular patterns (DAMPs), forming a vicious cycle that exacerbates tubular injury(Raup-Konsavage et al., 2018). In long-term chemotherapy models, intestinal injury has been demonstrated to result in LPS leakage and persistent low-grade endotoxemia, with some LPS serving as a continuous source of circulating NETs(F et al., 2023). Repeated low-dose cisplatin has also been shown to be a significant inducer of NETosis, a finding that is consistent with the observations reported in this study. In light of these findings, compromised physiological barriers appear to allow extensive NET infiltration into the renal parenchyma, resulting in direct damage to tubular cells and activation of macrophage inflammasomes. This process ultimately leads to the development of interstitial fibrosis.

Given that LPS represents a significant component of the outer membrane phospholipids of the majority of gram-negative bacteria in the gut and elicits a range of inflammatory immune responses in the host if it enters the circulation, we therefore measured LPS levels in serum and leakage of mouse intestines. Our results indicated that OPCs restored the serum LPS level and blocked cisplatin-induced intestinal leakage. In parallel, we also found OPCs up-regulated the intestinal tight junction proteins, suggest that OPCs inhibited LPS leakage via targeting the intestinal barrier. Tight junction proteins play essential roles in maintaining the structural integrity of the intestinal barrier. Their functions are carried out through transmembrane proteins such as claudin, Occludin, and tricellulin, as well as intracellular scaffolding proteins, including ZO-1, ZO-2, and ZO-3(Ghosh et al., 2021; Mouries et al., 2019). Consistent with this data, some studies demonstrated that OPCs also restored intestinal tight junction proteins, thereby inhibiting increased permeability in several systemic inflammation-related diseases, (González-Quilen et al., 2019; Xu et al., 2019).

We next aimed to understand how OPCs restored tight junction proteins. In rodents and humans, the composition and metabolism of the gut microbiota (ecological dysbiosis), characterized by increases in the overall diversity of the microbiota and the abundance of Bacteroidetes (including Muribaculaceae, Clostridia-UCG-014, Lachnospiraceae NK4A136, et. al)contribute to the integrity of tight junction. It is reported that the increase of Muribaculaceae and Clostridia-UCG-014 induce intestinal mucus secretion and enhance intestinal mucus barrier function(Q. Wang et al., 2023), which limit LPS levels in the bloodstream and ameliorate metabolic endotoxemia(Li et al., 2016). The Lachnospiraceae NK4A136 group also has a significant inverse correlation with intestinal permeability and plasma LPS levels and a positive correlation with colon length and Cldn1 expression. In agreement with these findings, our flora study demonstrated that the Muribaculaceae Akkermansiaceae, Lachnospiraceae_NK4A136, and Glostridia_UCG-014, exhibited a decline in response to RLDC, after that the OPCs subsequently recovered them, indicating that OPCs protected the intestinal mucosal barrier via increase of Bacteroidetes abundance. What’s more, the increase in abundance of Bacteroidetes decreases the concentrations of indophenol sulfate and p-cresol sulfate-based uremic toxins, which in turn improve renal function(Kieffer et al., 2016). The uremic toxins from CKD patients increase intestinal permeability, resulting in marked deficiencies in crucial protein components of epithelial tight junctions (Claudin-1, Occludin, and ZO-1). This may subsequently lead to bacterial and endotoxin translocation across the intestinal wall(Vaziri et al., 2013, 1985). Thus, it is reasonable to believe that OPCs-increased the above flora is required for intestinal integrity to inhibit LPS leakage. These findings further demonstrate that the gut and kidney act interdependently, amplifying damage to the organism through the gutLJkidney axis. OPCs protected the kidneys by restoring the physical and ecological structure of the gut, thereby disrupting the positive feedback pathway of the gutLJkidney axis.

Notably, in our study, we used BBR as a positive control drug to kill intestinal bacteria and found that BBR reduced intestinal inflammation and alleviated CKD, but this effect was not as effective as that of OPCs. It may be attributed to the function of OPCs in adjusting the gut microbiota while BBR only inhibited the growth of all bacteria (data not shown). Another reason could be the direct effect of OPCs being absorbed into the circulation. Previous studies conducted in our laboratory have demonstrated that OPCs inhibited NLRP3 inflammasome activation by scavenging ROS. It reduced CASP1 and IL-1β levels, thereby relieving gout pain(Liu et al., 2017). It has been reported that OPCs can reduce cisplatin-induced oxidative stress and inflammatory infiltration, alleviating acute nephrotoxicity(Sayed, 2009). Consistent with our data, we found OPCs decreased the levels of ROS, NETs, NLRP3, CASP1, IL18, and IFNγ and prevented renal collagen deposition and fibrosis formation (Figure 5). The activation of ROS plays a key role in NETosis(Amara et al., 2021), as well as in CKD. Thus, we supposed that OPC suppressed NETs and inflammatory responses by scavenging of ROS, ultimately alleviating CKD.

With respect to the roles of NETs in cisplatin-treated mice with AKI, it has been reported that targeting NETs can improve renal function(Mousset et al., 2023) and alleviate AKI, which is due to the key role of NETs in ischemia(Raup-Konsavage et al., 2018) and sepsis(Ni et al., 2021). However, the effects of NETs in CKD caused by RLDC and the related mechanisms in kidney injury remain obscure. Cisplatin is primarily cleared by the kidneys through glomerular filtration and tubular excretion, resulting in higher concentrations in the kidneys than in other organs; thus, renal accumulation is associated with a high incidence of acute and chronic nephrotoxicity. In general, the high affinity of high-dose cisplatin to DNA directly induces necrosis and endoplasmic reticulum stress-related apoptotic pathways, leading to cell death(Wei et al., 2007). In contrast to the nephrotoxic effects of high-dose cisplatin, low-dose cisplatin, which leads to accumulation of the drug in regeneratively repaired renal tubular epithelial cells, does not result in severe toxicity. However, upon the intestinal leakage, endotoxins entering the bloodstream combined with low concentrations of cisplatin yielded different results. Our cellular experiments imitated the in vivo microenvironment under the dual effects of endotoxins and cisplatin using L-LPS plus L-CIS. Surprisingly, L-LPS plus L-CIS could activate the production of NETs, whereas neither L-LPS nor L-CIS alone could unequivocally induce NETs formation (Figure 9). It indicated that low endotoxin state in circulation facilitated the effect of low-dose cisplatin on CKD. It induced NETs formation, maintaining a proinflammatory state, which ultimately led to chronic, irreparable nephropathy at the next cycle of cisplatin treatment(Yamashita et al., 2021). In this study, we found that NETs were indispensable in this chronic vicious cycle.

Elevated NETs in the kidneys have been shown to activate the NLRP3 inflammasome, resulting in the release of IL-18 and IFN-γ, which –induced tubulointerstitial inflammation and fibrosis. Consistent with this view, we found that PAD4 knockout inhibited NLRP3 expression, accompanied by decreases of CASP1, IL18, and IFNγ expression, in the RLDC model (Figure 3). Interestingly, OPCs decreased the expressions of these proteins and improved the renal function while NLRP3 inhibitors and IL-18 binding proteins did not improve renal function, suggesting that the NETs-NLRP3-CASP1-IL18-IFNγ pathway was involved in the progression of cisplatin-induced renal fibrosis but not crucial for CKD. Additionally, NETs provide a scaffold for the exposure of functional TFs in thrombus formation(Stakos et al., 2015; von Brühl et al., 2012) and concurrently, neutrophils secrete the protease MMP9, which contributes to local ischemiaLJhypoxia (Carmona-Rivera et al., 2015). In a manner analogous to its function in renal development, MMP-9 inhibits apoptosis in the AKI model, which is renoprotective(Bengatta et al., 2009). In contrast to fibrin-degrading activity during AKI, MMP9 plays a negative role during the period of AKI to CKD progression and promotes EMT with fibrosis in a CKD model(Wang et al., 2019). Our previous data demonstrated that neutrophil infiltration was accompanied by elevated TF and MMP9 in a mouse model of chemotherapeutic enteritis and endotoxin(Li et al., 2021; Yu et al., 2020). In this study, we found that the levels of NETs were elevated, accompanied by elevated TF and MMP9 and diminished TFPI during RLDC-induced CKD (Figure 4). Furthermore, OPCs reversed the effect of RLDC on CKD, revealing that OPCs inhibited the formation of microthrombi by inhibiting NETs formation. Thus, we believe that OPCs may act as a protective agent against cisplatin nephropathy by inhibiting NETs-mediated inflammation and ameliorating ischemia and hypoxia (Figure 10).

**Figure 9.**
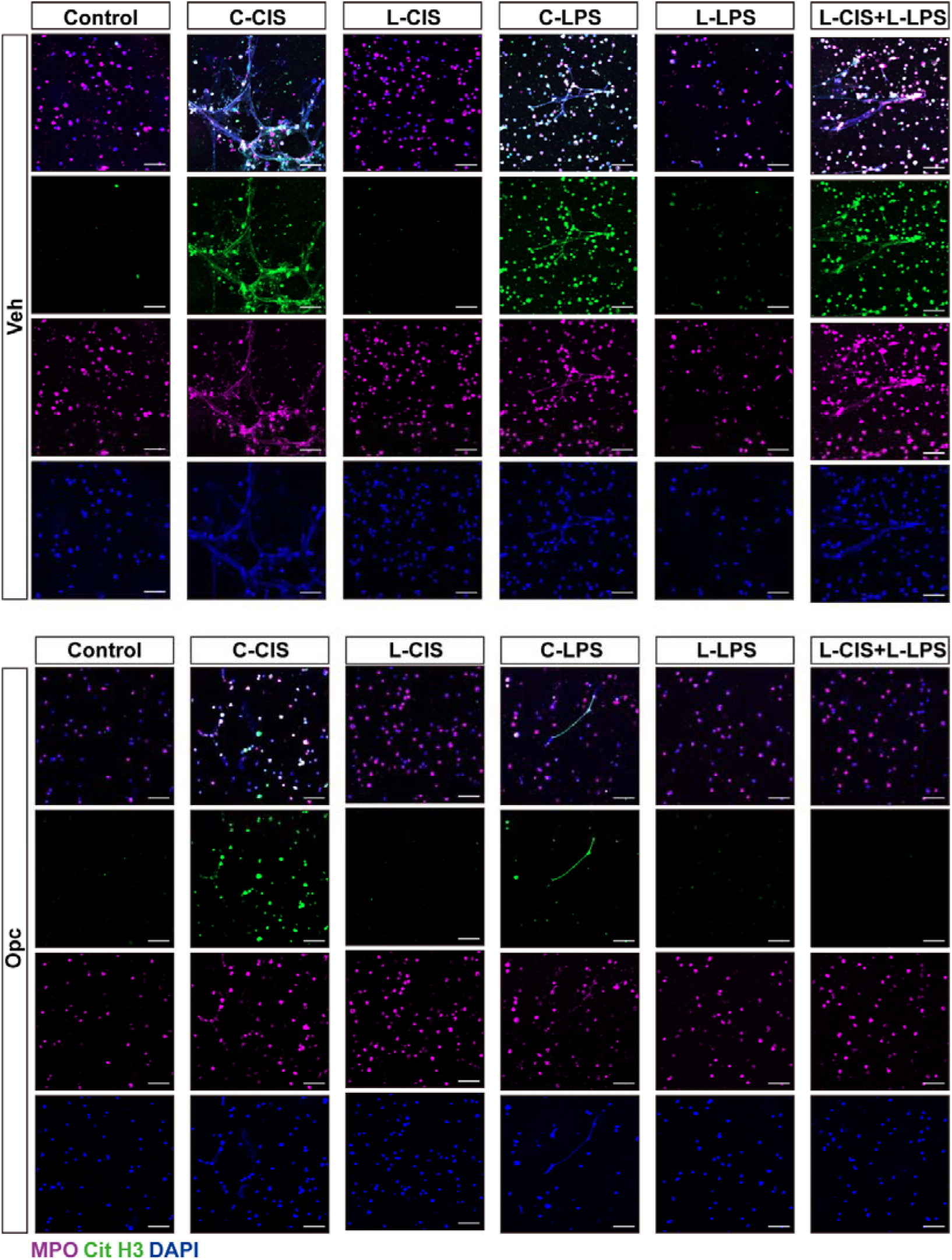
OPCs inhibits the cisplatin– and LPS-induced release of NETs from neutrophils. **A, B.** Immunostaining of mouse neutrophils cultured as indicated. Anti-MPO (purple), anti-Cit H3 (green), and DAPI (blue) staining was used to assess NETs formation. Scale bar = 100 μm (*n* = 3).

**Figure 10.**
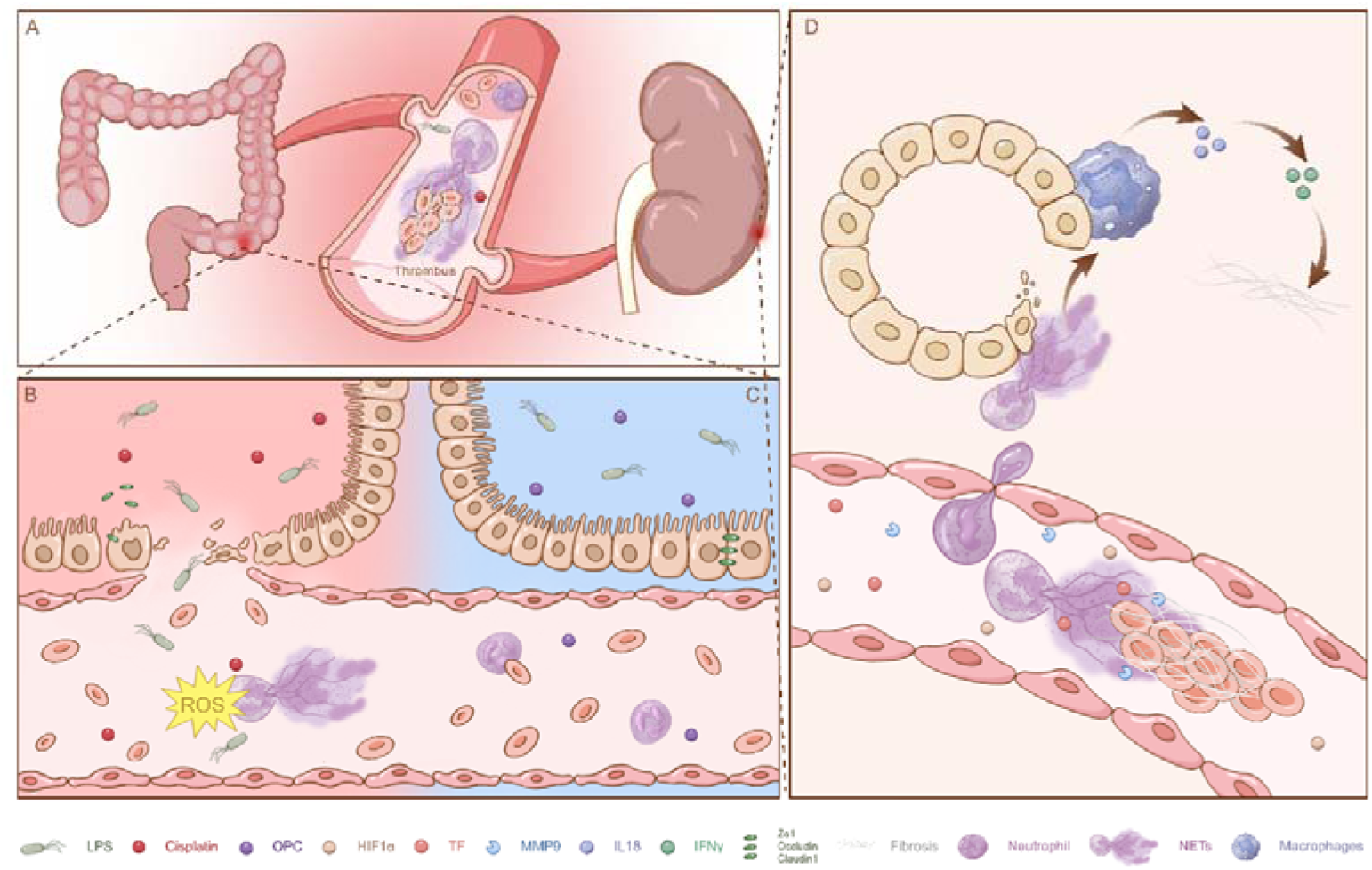
Schematic illustration showing that RLDC-induced NETs promote the development of CKD by disrupting the gut barrier and the therapeutic role of OPCs. **A.** The combination of cisplatin and intestine-derived LPS induces NETs formation, leading to CKD via the gutLJkidney axis. **B.** Cisplatin-induced gut barrier dysfunction facilitates NETosis caused by both LPS and cisplatin, which disturb microcirculation. **C.** The inhibition of NETosis by OPCs is attributed to its anti-inflammatory and antioxidant activities and ability to maintain a balanced intestinal flora. **D.** NETs induce local ischemia and fibrosis, which are involved in the pathogenesis of kidney damage.

In the kidney, proximal tubular cells are the most susceptible to hypoxic stimuli, and the extent of tubular injury is a crucial determinant of the prognosis of renal disease. Hypoxia-promoted coagulation and thrombotic events play important roles in the pathogenesis of CKD(Oe and Takahashi, 2022). Indeed, NETs can form large aggregates that may even be large enough to block small blood vessels without coagulation activation (Jiménez-Alcázar et al., 2017), further exacerbating the susceptibility of the kidney to hypoxia. The high oxygen consumption of the renal tubules, coupled with the relatively low blood circulation in the proximal tubules of the kidney, exacerbates renal injury further. During the period of renal repair, an increase in aerobic glycolytic oxygen consumption leads to relative hypoxia during the period of renal repair, causing a vicious cycle involved in renal injury(Liu et al., 2018). We found Neutrophils treated with RLDC were more prone to form NETs, and the damage to renal tubular epithelial cells was much greater than that of cisplatin alone (Supplementary Figure 5-7), indicating the key role of NETs in RLDC-induced CKD. Futher, our study revealed that the increase in the renal hypoxia-inducible factor HIF1α was proportional to the increase in the proliferative marker protein Ki67 after RLDC induction, which suggested that hypoxia and repair formed a malignant cyclic process in CKD, whereas the absence of NETs-associated pathological emboli after PAD4 knockout impeded the malignant positive feedback process at the source. In conclusion, the advantage of NETs is that they facilitate bidirectional communication between the immune response and coagulation, thereby enhancing the efficacy of the two major host protection systems, hemostasis, and innate immunity.

In conclusion, prolonged chemotherapy leads to impaired intestinal barrier function, allowing LPS to enter the circulation and induce NETs, which are eventually deposited in the kidney, leading to inflammatory fibrosis and necrosis due to ischemia and hypoxia, which promotes the progression of CKD in the kidney. What’s more, OPCs exert anti-inflammatory, antibacterial, and antioxidant effects, thereby preventing the progression of CKD.

## Materials and methods

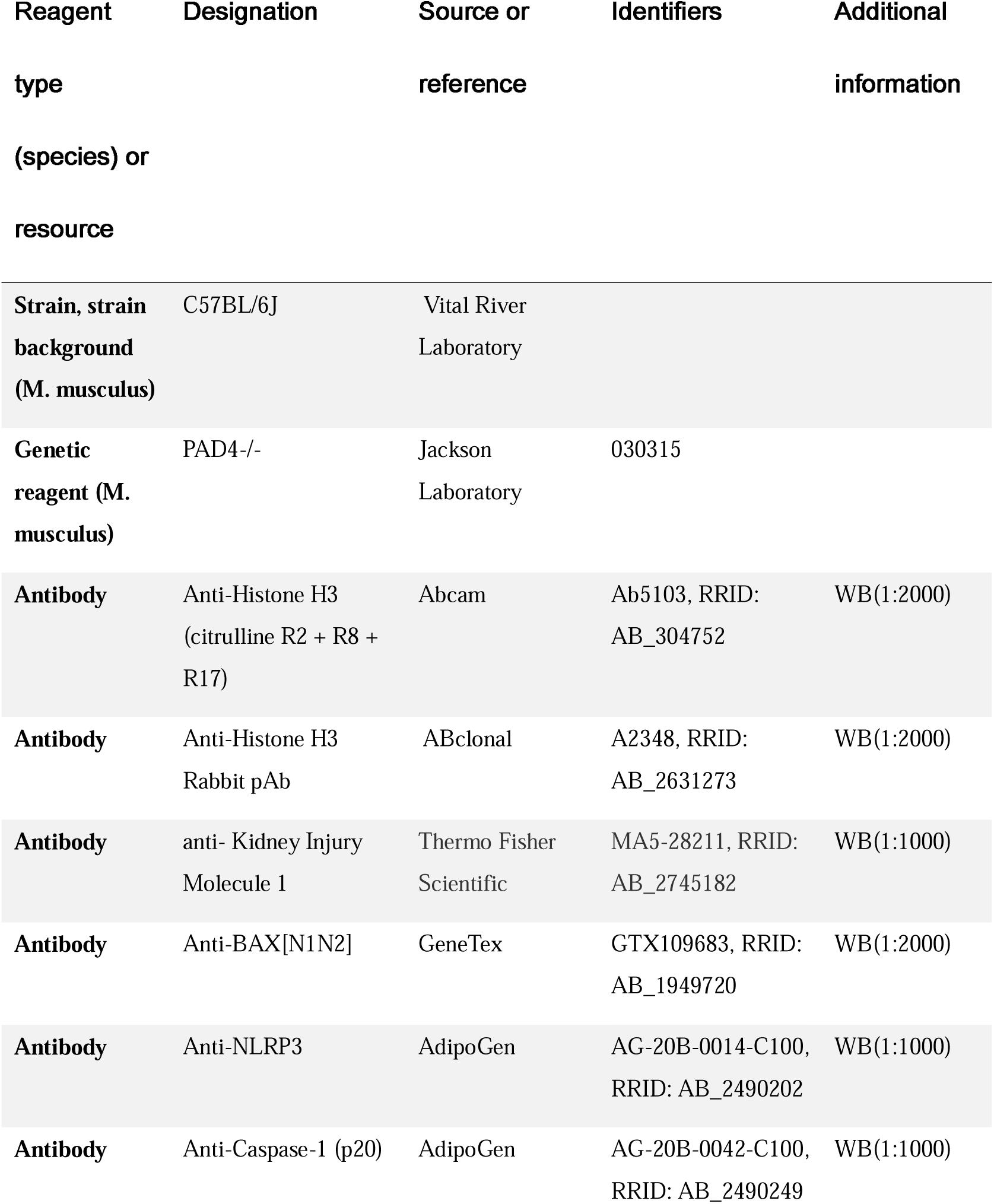
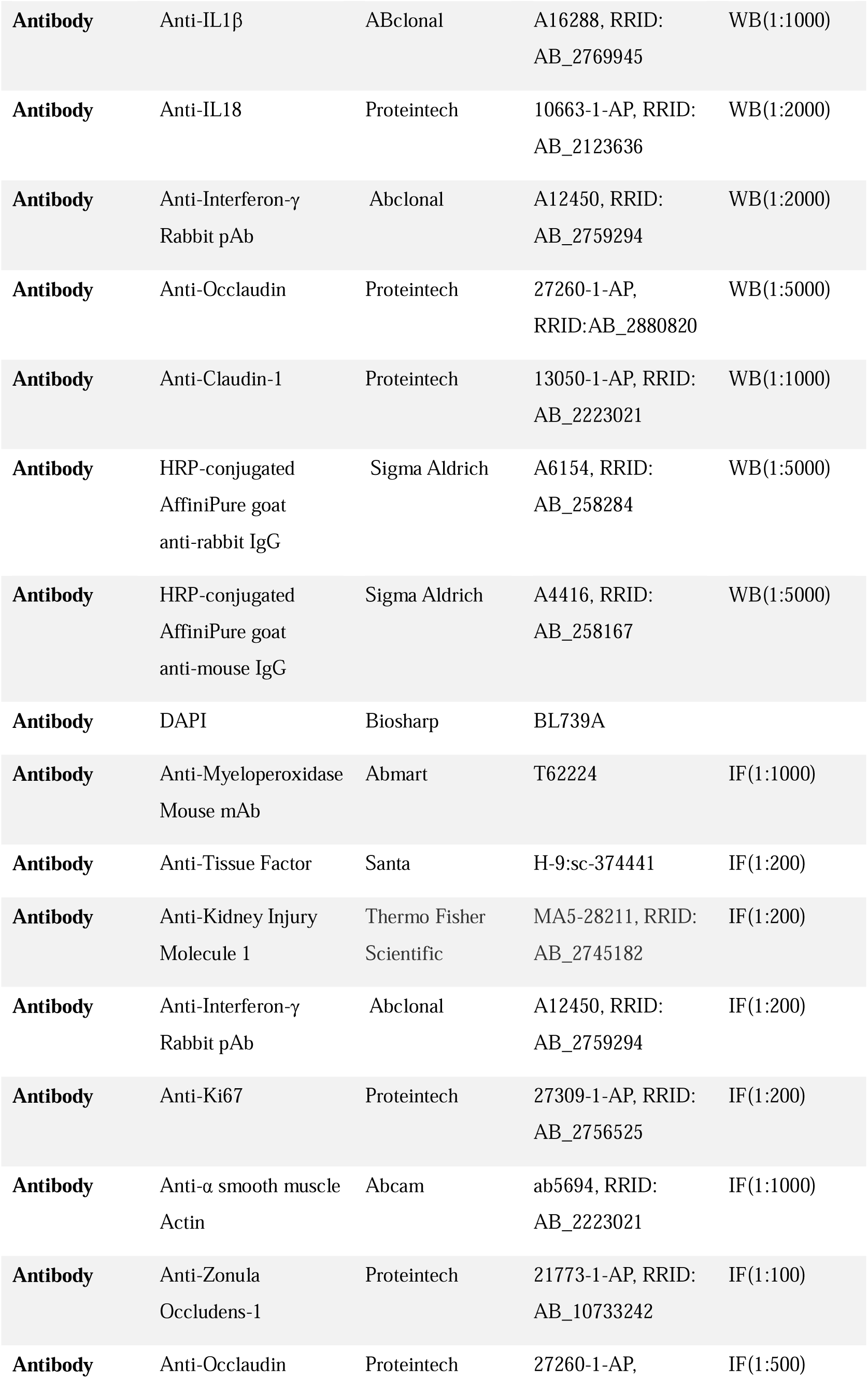
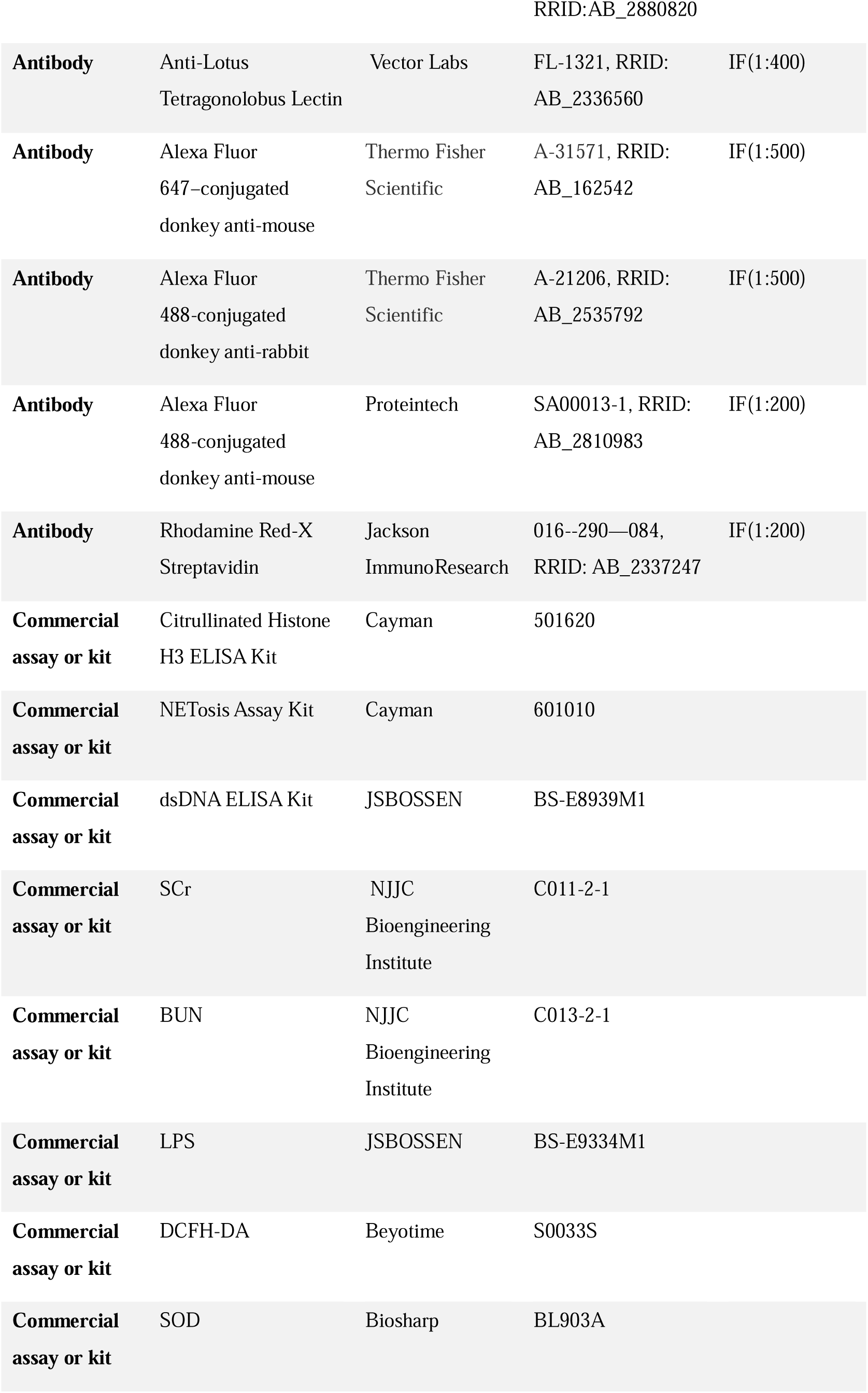
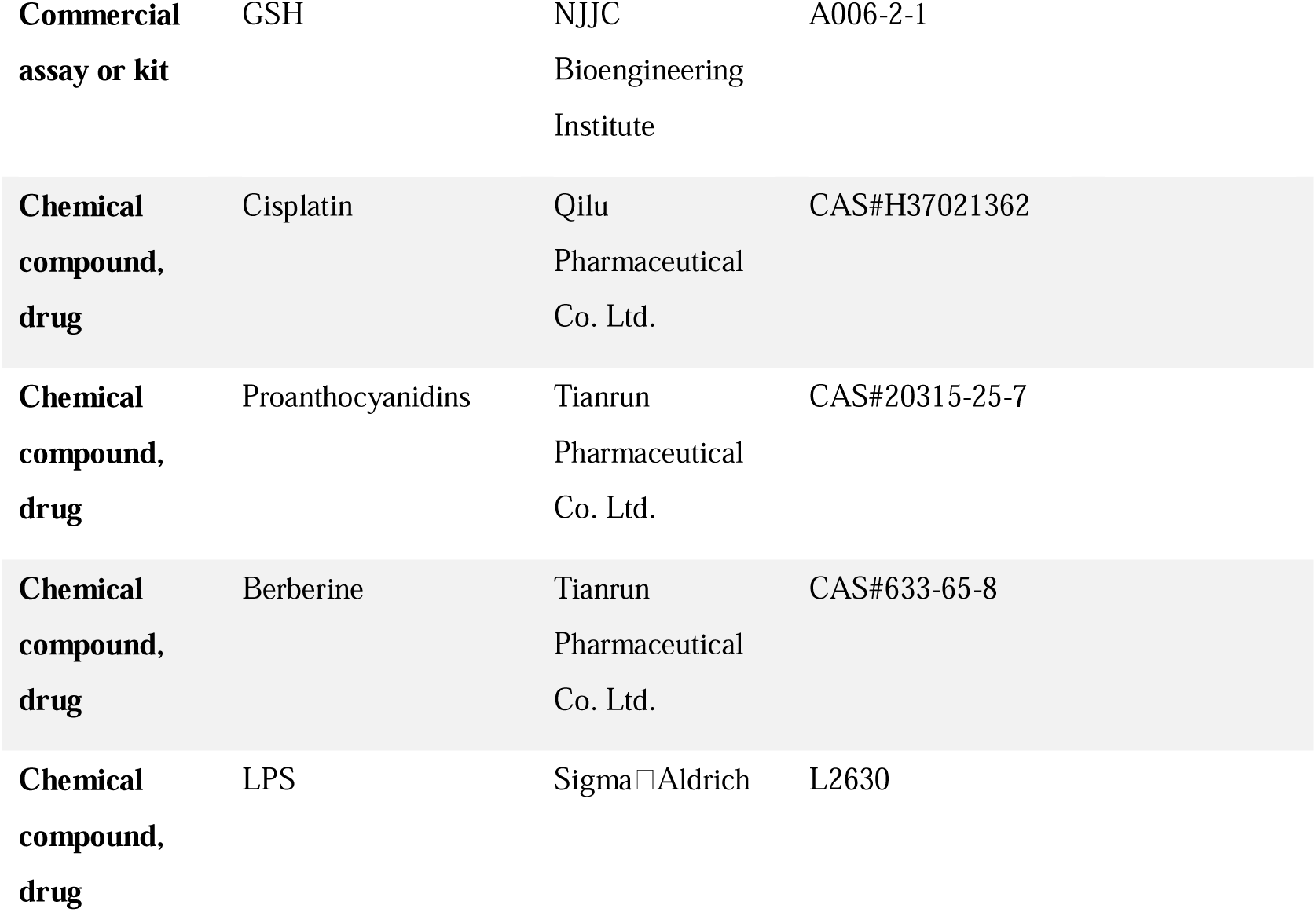
Key resources table.

### Mouse model of RLDC and OPCs treatment

Adult male C57BL/6J mice (20–25 g, wild type) were obtained from the Animal Core Facility of Nanjing Medical University, Nanjing, China. Pad4−/− mice on a C57BL/6J background were purchased from The Jackson Laboratory. The animals were housed five to six per cage under pathogen-free conditions with soft bedding at a controlled temperature (22°C±2°C) and photoperiod (12:12-hour lightLJdark cycle). Prior to the commencement of the experiment, the animals were permitted to acclimatize to the experimental conditions for a minimum of two days. The animals were matched for age and weight in each set of experiments. Eight-week-old male C57BL/6J background mice and PAD4−/− mice were intraperitoneally injected with 7 mg/kg cisplatin (CIS, H37021362, Qilu Pharmaceutical Co. Ltd.) or 0.2 ml of saline via the intraperitoneal route on a weekly basis for a period of four weeks(Sears and Siskind, 2021).

To assess the therapeutic effects of natural medicines, mice were administered OPCs. OPCs (100 mg/kg; Tianrun Pharmaceutical Co. Ltd.) was dissolved in 0.5% carboxymethyl cellulose sodium (A501427-0250; Sangon Biotech) and was given by gavage three times a week for four weeks, commencing with the initial cisplatin injection. Berberine (100 mg/kg; Tianrun Pharmaceutical Co. Ltd.) was administered via gastric lavage three times a week for four weeks, starting with the initial cisplatin injection as a positive control for treating the gut microbiota. The dosages of OPCs and BBR used in the experiments were previously described and were based on the results of preliminary experiments(Allameh et al., 2020; Gao et al., 2023; Gholampour et al., 2022). The purity of the OPCs was greater than 95%. The OPCs contained 1.1% monomeric procyanidins, 34.2% dimeric procyanidins, 24.9% trimeric procyanidins, 6.7% tetrameric procyanidins (66.9% oligomeric procyanidins in total), and 33.1% polymeric procyanidins.

### Histological staining

Kidney tissues were fixed with 4% paraformaldehyde, embedded in paraffin, and sectioned at 4 μm. Renal tissues were subjected to hematoxylin and eosin (HE) staining. The extent of tubular damage was quantified at 40× magnification, with a total of 200 cortical tubules examined and calculated in accordance with previously described methods (Li et al., 2018; Tan et al., 2023). For periodic acid-Schiff (PAS) staining, the lumen of the tube containing glycogen components was stained red–purple. Masson’s trichrome staining was employed to assess the presence and distribution of collagen fibrils within renal tissues. For quantification, 6 positive collage-stained fields (40× magnification) were randomly selected from each section and analyzed via ImageJ(Fu et al., 2019).

### Colon Tissue Processing

The distal colon was removed and washed with a physiological saline solution. Segments of approximately 0.5 cm in length were then cut. A proportion of the colon samples were fixed in 4% paraformaldehyde solution, prepared in 0.1 M phosphate-buffered saline (PBS) with a pH of 7.4, and then embedded in paraffin wax for morphological analysis. A proportion of the colon samples were fixed in 4% paraformaldehyde solution, prepared in 0.1 M PBS at a pH of 7.4, and then embedded in paraffin wax for morphological analysis. A minimum of 40 villus lengths were quantified on selected HE-stained sections via ImageJ software(Zhan et al., 2023).

### Immunofluorescence

After administering anesthetic agents to the mice, the kidneys and intestinal tissues were excised via a transcardiac infusion of saline, fixed in 4% paraformaldehyde, and the embedded blocks were sectioned into 4 μm thick pieces. These samples were then blocked for one hour with 1% normal donkey serum (017--000--121, Jackson ImmunoResearch, RRID: AB_2337258) and 0.1% Triton-X in phosphate buffer solution (PBS). For immunofluorescence analysis, tissue sections were subjected to incubation with primary antibodies against Cit H3 (ab281584, Abcam), MPO (T62224, Abmart), TF (H-9:sc-374441, Santa), KIM-1 (MA5-28211, Thermo Fisher Scientific, RRID: AB_2745182), Ki67 (27309-1-AP, Proteintech, RRID: AB_2756525), IFN-γ (A12450, ABclonal, RRID: AB_2759294), α-SMA (ab5694, Abcam, RRID: AB_2223021), ZO-1 (21773-1-AP, Proteintech, RRID: AB_10733242) and Occlaudin (27260-1-AP, Proteintech, RRID: AB_2880820) overnight at 4°C. The proximal tubules were labeled with LTL (FL-1321, Vector Labs, RRID: AB_2336560), and the nuclei were counterstained with DAPI. The secondary antibodies used were as follows: Alexa Fluor 647–conjugated donkey anti-mouse (A-31571, Thermo Fisher Scientific, RRID: AB_162542), Alexa Fluor 488-conjugated donkey anti-rabbit (A-21206, Thermo Fisher Scientific, RRID: AB_2535792), Alexa Fluor 488-conjugated donkey anti-mouse (SA00013-1, Proteintech, RRID: AB_2810983), and Rhodamine Red-X Streptavidin (016--290--084, Jackson ImmunoResearch, RRID: AB_2337247). Following three washes with PBS, the samples were examined under a fluorescence microscope (Leica DM2500) to ascertain the morphological details of the immunofluorescence staining. The examination was conducted in a blinded manner.

### Neutrophil isolation

The extraction of bone-marrow-derived neutrophils is achieved through a gradient density centrifugation technique. After the mice were euthanized, the tibias and femurs were isolated via aseptic techniques. The ends of the bones were then cut after the muscle tissue was removed to expose the marrow cavity. A 5 ml syringe was filled with RPMI-1640 (KGM31800N-500, Keygen Biotech), 10% FBS (04-001-1ACS, Biological Industries), and 2 mmol/L EDTA (E9884, SigmaLJAldrich) to flush the bone marrow cells into a 50 ml centrifuge tube. After centrifugation at 1200 × g for 5 min at 4°C, the cells were resuspended in 3 ml of sodium chloride. Then the supernatant was added to the upper layer of 9 ml of Histopaque-1077 (density, 1.077 × g/mL, 10771, SigmaLJAldrich) and centrifuged at 2000 × g for 20 min at 4°C without braking. The supernatant was discarded, and the cells were resuspended in 5 ml of sodium chloride physiological solution. Neutrophils were collected at the junction of tissue water and sodium chloride physiological solution after the cells were placed in the upper layer of 10 ml of Histopaque-1119 (density, 1.119 g/mL, 11191, SigmaLJAldrich) and centrifuged for 20 min at 4°C and 2000 × g without braking. The collected neutrophils were washed twice with RPMI 1640 supplemented with 10% FBS and 1% penicillin/streptomycin (KGY0023, Keygen Biotech) and centrifuged at 1400 rpm for 7 min at 4°C.

In order to isolate peripheral blood neutrophils from mice, it is first necessary to employ the Mouse Peripheral Blood Neutrophil Isolation Solution Kit (P9201, Solarbio) to mix fresh anticoagulated blood with PBS and erythrocyte sedimentation medium in a 1:1:1 ratio, in accordance with the instructions. Allow the mixture to stand at room temperature for 30 minutes to facilitate removing red blood cells. Subsequently, the upper layer of the mixture was collected and mixed with Reagent A and Reagent C. The mixture was then subjected to a centrifugal process at 1000 × g for 30 minutes at ambient temperature. Aspirate the contents of the lower milky-white ring, add cell wash solution, centrifuge at 250g for 10 minutes, discard the supernatant, and resuspend the cells.

### Neutrophil treatment

To assess the promotion of NETs formation by LPS versus cisplatin and the inhibition of NETs formation by OPCs, neutrophils were incubated with C-LPS (1 mg/ml, L2630, SigmaLJAldrich), C-CIS (3 μg/ml, HY-17394, MedChemExpress) or L-LPS (10 ng/ml, L2630, SigmaLJAldrich), or L-CIS (0.15 μg/ml, HY-17394, MedChemExpress) for 4 hours, and OPCs (1 µg/ml, Zelang Pharmaceutical Co. Ltd.) or RPMI1640 was added to the medium 1 hour before LPS or cisplatin.

### NETs isolation

Extracted neutrophils were then stimulated with C-CIS (3 μg/ml, HY-17394, MedChemExpress) in RPMI 1640 medium for a period of 24 hours. Following the meticulous removal of the upper layer, the NETs that had adhered to the bottom of the well were gently resuspended in PBS. Subsequently, the mixture was subjected to a centrifugation process at 1000g for a duration of 10 minutes at a temperature of 4°C. This procedure was undertaken to achieve the formation of a NETs pellet. The pellet should be dissolved in 100 μl of PBS, and the quantity of purified cell-free NETs subsequently quantified using a microplate reader in conjunction with a NETs detection kit, for use in further experiments.

### Renal tubular epithelial cells treatment

Following the euthanasia of the mice, the renal cortex tissue must be collected, minced, and resuspended in 10 mL Hank’s Balanced Salt Solution (HBSS). Five millilitres of 0.1% type II collagenase should then be added, and the mixture should be incubated at 37°C in a water bath for 30 minutes, with frequent inverting to ensure thorough mixing. The suspension should then be filtered through a cell strainer (70 μm, Beyotime), after which the glomerular cells should be removed by grinding and passing the suspension through a 40 μm cell strainer. The suspension should then be subjected to a centrifugal process at a speed of 1000 revolutions per minute for three minutes. The tubular cells were collected post-ultracentrifugation in a density gradient. Following a thorough wash with PBS, the pellet should be resuspended in Dulbecco’s Modified Eagle Medium (DMEM)/F12 (1:1) medium, with the addition of 10% Fetal Bovine Serum (FBS), and left to incubate for a period of 4-5 days. The treatment of choice is the administration of extracted NETs at a concentration of 500 ng/ml, or alternatively, a concurrent treatment involving DNase I, NAC, and OPCs for a duration of 24 hours.

### In vitro NETs assay

For immunofluorescence staining, freshly isolated polymorphonuclear neutrophils (PMNs) were seeded on poly-D-lysine-coated coverslips, and their adherence was permitted. After NETs production was induced, the cells were fixed for 15 minutes with 4% paraformaldehyde (PFA) and blocked with 1% BSA and 0.3% Triton X-100 in PBS for 30 minutes. Then, anti-Cit H3 and anti-MPO (ab90810, Abcam) primary antibodies were used overnight at 4°C. After three washes, Alexa Fluor 488-conjugated donkey anti-rabbit (A-21207, Invitrogen), Alexa Fluor 647-conjugated donkey anti-mouse (A-31571, Invitrogen), and DAPI (BL739A, Biosharp) were added for 2 hours at room temperature. NETs formation was visualized via fluorescence microscopy (Leica DM2500).

### Quantification of NETs

Mouse plasma was collected from whole blood by centrifugation at 3,000 rpm for 5 minutes. The quantification of NETs in plasma was conducted in accordance with the Citrullinated Histone H3 ELISA Kit (501620, Cayman), NETosis Assay Kit (601010, Cayman), and dsDNA ELISA Kit (BS-E8939M1, JSBOSSEN). The absorbance was quantified with a microplate reader (Multiskan FC, Thermo Fisher).

### Renal function

In summary, blood samples were taken from the infraorbital region for anticoagulation and then centrifuged at room temperature to collect the serum. For Scr (C011-2-1, NJJC Bioengineering Institute), the reaction was conducted at 37°C for 5 min, and the absorbance at 546 nm was recorded at the end of the response. For BUN (C013-2-1, NJJC Bioengineering Institute), the sample was added to the preheated reaction mixture at 37°C, and the absorbance at 640 nm was monitored after 10 min of reaction. BUN and Scr levels (mg/dl) were calculated according to the assay kit.

### LPS assay

All materials used for both sample preparation and testing were pyrogen-free. The lipopolysaccharide (LPS) concentration in the serum was quantified via a chromogenic endotoxin assay (BS-E9334M1, JSBOSSEN) based on a Limulus amebocyte extract. The samples were subjected to centrifugation at 3000 rpm for a period of 10 minutes to separate the supernatant. The endotoxin concentration was expressed in endotoxin units per milliliter (EU/mL). All the data were obtained from standard curves.

### Measurement of ROS

Neutrophil collection was performed subsequent to the induction of NETs production, and DCFH-DA (S0033S, Beyotime) was used to quantify reactive oxygen species (ROS) as a fluorescence probe. Upon cell formation, DCFH is oxidized by intracellular ROS and converted to DCF. Consequently, the observed fluorescence signal was found to be proportional to the production of ROS. A laser confocal microscope was used to observe the staining results, and the fluorescence intensity was measured at an excitation wavelength of 488 nm and an emission wavelength of 525 nm with a fluorescence enzyme marker (Cytation).

### Flow cytometry

In order to assess neutrophil peroxidase levels across different groups, samples were treated with the ROS assay kit (S0033S, Beyotime) and analysed via flow cytometry. The collected neutrophils must then be resuspended in DCFH-DA, which has been diluted with serum-free medium. The mixture should then be subjected to an incubation process at a temperature of 37°C within a cell culture incubator for a period of 20 minutes, with the mixture being inverted at intervals of between 3 and 5 minutes. The treated neutrophils are then washed thrice with serum-free cell culture medium, after which the resuspension is analysed using flow cytometry (Miltenyi MACSQuant Analyser 10). For each sample, a total of 10,000 cells were collected, and the ratio of DCFH-DA positive cells was analyzed through flow cytometry. The data were analyzed using FlowJo statistical software (v10.62).

To assess the resistance of peripheral blood neutrophils across different groups, the Annexin V-FITC apoptosis detection kit (C1062M, Beyotime) was employed. Extracted neutrophils were cultured in 6-well plates and stimulated with L-CIS (0.15 μg/ml, HY-17394, MedChemExpress) for 24 hours. Following the incubation period, the upper layer of the mixture was discarded, and the cells were washed with PBS. Subsequently, 100,000 cells were resuspended in 195 μl of Annexin V-FITC binding buffer. The next step is to add 5 μl of Annexin V-FITC and 10 μl of propidium iodide. These should be added gently, and the mixture should then be incubated at room temperature in the dark for 20 minutes. The cells were then washed with warm PBS, and subsequently transferred from the culture plate using cold PBS containing 1% foetal bovine serum for the purpose of flow cytometric analysis (Miltenyi MACSQuant Analyser 10). The subsequent analysis of the data was conducted utilising the FlowJo software version 10.62.

### Measurement of SOD and GSH

Mouse blood was collected in an anticoagulation tube, mixed upside down, and centrifuged at 600 × g for 10 min at 4°C, and the supernatant was added to the working solution and incubated at 37°C for 30 min. The absorbance at 450 nm was then measured to detect the inhibition rate of the SOD enzyme in the blood (BL903A, Biosharp). For GSH (A006-2-1, NJJC Bioengineering Institute), 0.05 ml of 10-fold diluted heparin anticoagulated blood and 0.2 ml of working reagent were centrifuged at 3500 rpm for 10 min. The supernatant was removed, the subsequent working solution was added, the mixture was oscillated, the mixture was mixed for 5 min, and the absorbance at 405 nm was detected.

### Western blotting

Following the euthanasia of the animals, their kidneys and colon tissues were rapidly excised and homogenized in RIPA lysis buffer after protein concentration was measured with a Pierce™ BCA protein assay kit (23225, Thermo Fisher Scientific). Equal amounts of protein (40 μg for each tissue lysate) were introduced, separated via SDSLJPAGE under reducing conditions, and subsequently transferred to PVDF membranes (0000206738, Millipore Corp.) for the standard procedure of immunoblot analysis. The membranes were blocked with 5% bovine serum albumin for 1 h at room temperature and probed with primary antibodies, including anti-histone H3 (A2348, ABclonal, RRID: AB_2631273), anti-citrulline histone H3 (ab5103, Abcam, RRID: AB_304752), anti-KIM-1 (MA5-28211, Thermo Fisher Scientific, RRID: AB_2745182), anti-BAX (GTX109683, GeneTex, RRID: AB_1949720), anti-NLRP3 (AG-20B-0014-C100, AdipoGen, RRID: AB_2490202), anti-caspase-1 (AG-20B-0042-C100, AdipoGen, RRID: AB_2490249), anti-IL1β (A16288, ABclonal, RRID: AB_2769945), anti-IL18 (10663-1-AP, Proteintech, RRID: AB_2123636), anti-IFNγ (A12450, ABclonal, RRID: AB_2759294), anti-Occlaudin(27260-1-AP, Proteintech, RRID:AB_2880820), and anti-Claudin-1 (13050-1-AP, Proteintech, RRID: AB_2079881).

The following secondary antibodies were used: HRP-conjugated AffiniPure goat anti-rabbit IgG (A6154, SigmaLJAldrich, RRID: AB_258284) and HRP-conjugated AffiniPure goat anti-mouse IgG (A4416, SigmaLJAldrich, RRID: AB_258167). Data were acquired with a molecular imager ChemiDoc system (ChemiDoc XRS+, Bio-Rad) and analyzed with ImageJ.

### Lower limb blood flow measurement

Lower limb blood flow was quantified via laser Doppler flowmetry (LDF). In particular, a computer-controlled optical scanner was employed to direct a low-power laser beam onto the exposed lower limb in a controlled fashion. Concurrently, the scanner head was positioned parallel to the exposed lower limb at a distance of approximately 20 cm. A colored image on the video monitor subsequently indicates the relative perfusion level in question. The values of blood flow are recorded and assessed by a Moor FLPIR view V40 program (Gene & I Scientific. Ltd), which has been designed for this purpose.

### Small animal imaging

Before autopsy, all the animals were fasted for 12 h and orally administered fluorescein isothiocyanate (FITC) dextran (60842-46-8, SigmaLJAldrich). After a four-hour incubation period, the distribution of fluorescein isothiocyanate (FITC) dextran in the mice was observed via a small animal imaging system (IVIS Spectrum, PerkinElmer). The peripheral blood of the mice was collected and centrifuged at 3000 r/min, 4°C, and 10 min, after which 100 μl of serum was obtained from the supernatant and added to a 96-well plate. The fluorescence intensity of the serum was quantified via a fluorescence microplate reader with an excitation wavelength of 480 nm and an emission wavelength of 520 nm.

### 16S rDNA Sequencing Analysis

Stool samples were stored at 80°C until analysis. Total stool DNA was extracted and isolated from the fecal pellets. A set of primers was designed to amplify a specific region, the 16S V3-V4 region, and a 420 bp fragment was successfully amplified. The fragment was then subjected to splicing via the Illumina NovaSeq 6000 platform, resulting in 2×250 bp paired-end data. These data were then subjected to splicing, which enabled the generation of a more extended sequence, thus facilitating 16S analysis. DADA2 was employed for the removal of low-quality sequences and chimeras, as well as for the generation of characteristic sequences for QIIME2 (https://forum.qiime2.org/t/qiime2-chinese-manual/838) for clustering operational taxonomic units (OTUs), diversity analysis, difference analysis, correlation analysis, and function prediction analysis. The default option for 16S rRNA genes is to use the Silva 138 rRNA database(Edgar, 2013). The proportion of sequences at different taxonomic levels for each sample was calculated on the basis of the abundance and annotation information of the ASVs. This was done to assess the species annotation resolution of the sample and the species complexity of the sample.

### Statistical analysis

The data from individual experiments are expressed as the mean ± standard error of the mean (SEM). To ascertain whether there were any significant differences between the groups, Student’s *t* test or ANOVA was employed, with a *p* value of less than 0.05 deemed to indicate statistical significance. The statistical analyses were conducted via Prism 9 software.

## Acknowledgments

We acknowledge all the authors who participated in this study. The authors thank the Center for Scientific Research of Nanjing Medical University and Shandong First Medical University for valuable help in our experiment.

## Additional information

### Funding

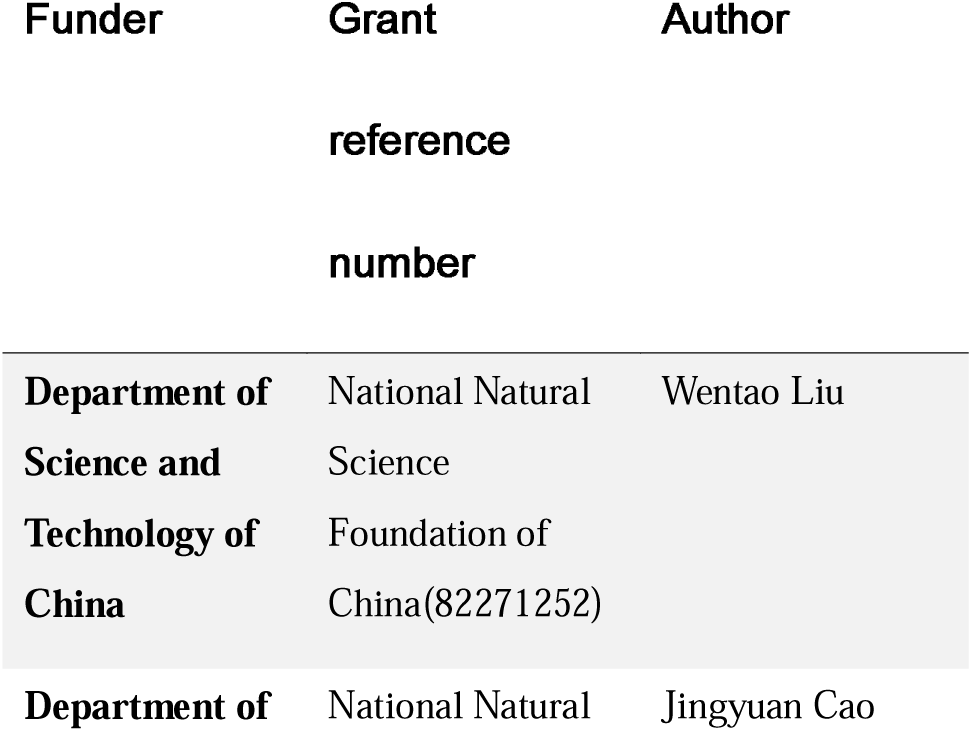
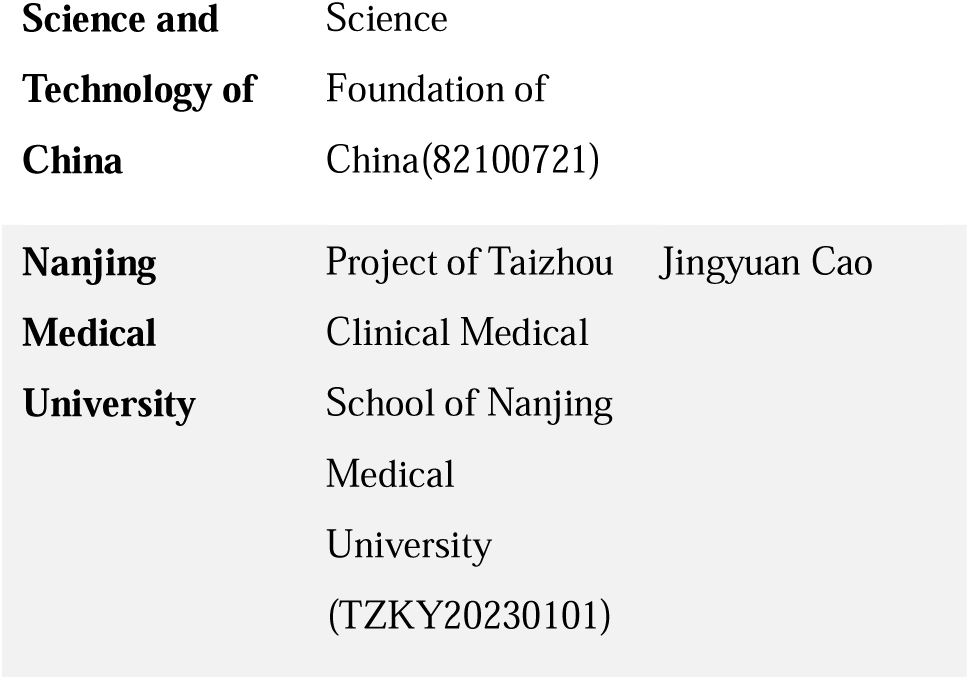

The funders did not have any role in the study design, data collection, data analyses, interpretation, or writing of the report.

### Author contributions

Yaqi Luan, Weiwei He and Kunmao Jiang designed and supervised the research. L., Shenghui Qiu, Lan Jin, Ying Huang and Xinrui Mao performed the experiments. Lai Jin and Wentao Liu wrote the original draft. Lai Jin, Jingyuan Cao and Rong Wang verified the data and edited the final draft. All the authors read and approved the final version of the manuscript, ensuring that this was the case.

### Ethics

All procedures were performed in strict accordance with the regulations of the ethics committee of the International Association for the Study of Pain and the Guide for the Care and Use of Laboratory Animals (The Ministry of Science and Technology of China, 2006). All animal experiments were approved by the Nanjing Medical University Animal Care and Use Committee (No. IACUC: 2305042) and were designed to minimize suffering and the number of animals used. All animal experiments complied with the ARRIVE (Animal Research: Reporting of In Vivo Experiments) guidelines (https://arriveguidelines.org).

### Data sharing statement

All data associated with this study are presented in the paper or the Supplementary Materials. Any additional information required to reanalyze the data reported in this work paper is available from the lead contact upon request.

## Figure and legends

**Supplement 1.**
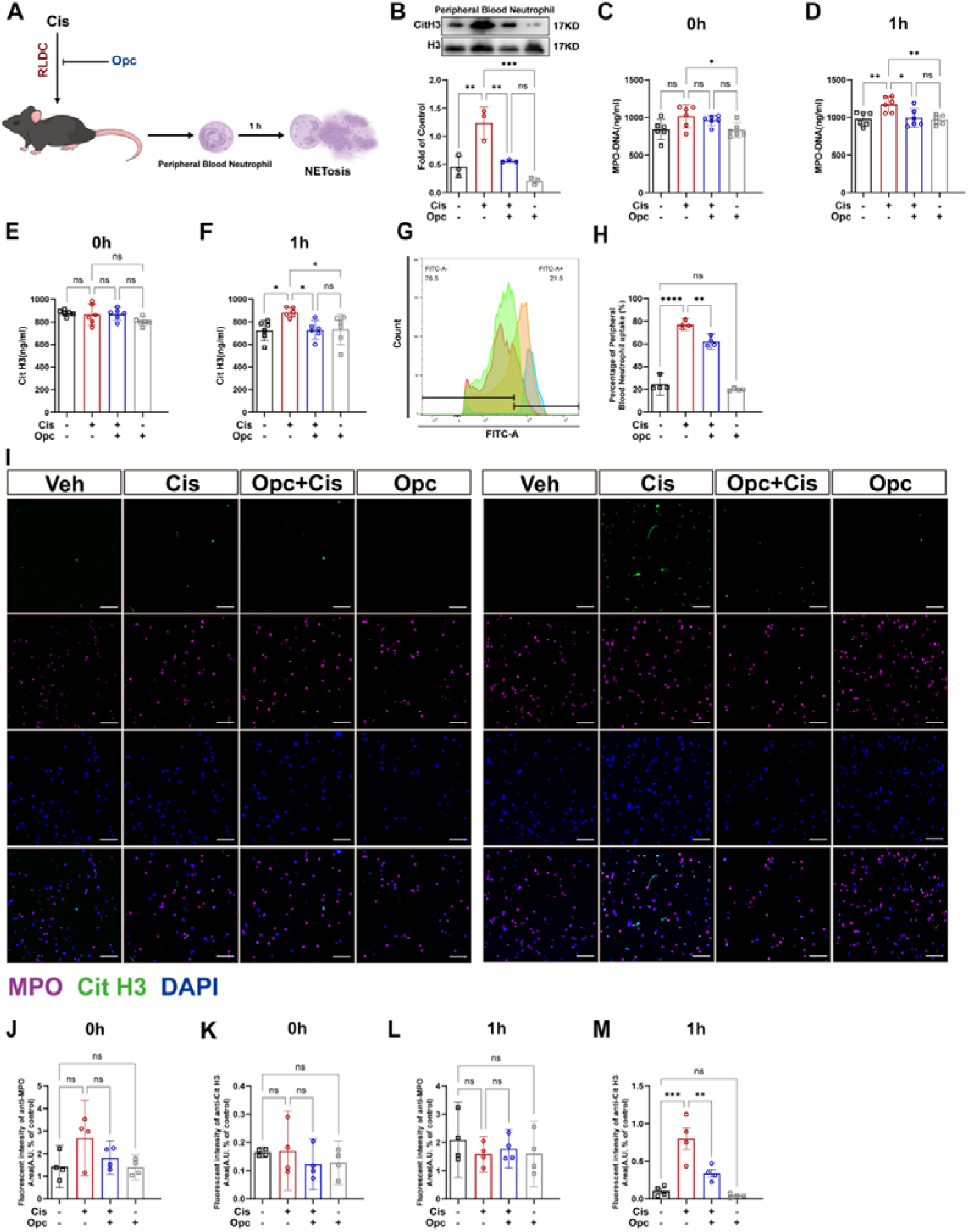
OPCs alleviate RLDC-induced NETosis. **A**. The scheme illustrating neutrophil extraction from peripheral blood of mice in different groups. **B.** Western blotting of Cit H3 protein levels in peripheral blood neutrophils. **C-F.** ELISA assays of MPO and Cit H3 in peripheral blood neutrophils at 0 hour or 1 hour post-cell extraction. **G-H.** Flow cytometry is used to assess the active state of reactive oxygen species in peripheral blood neutrophils. **I.** Immunofluorescence of MPO and Cit H3 in peripheral blood neutrophils. Anti-MPO (purple), anti-Cit H3 (green), and DAPI (blue) staining are used to assess NETs formation 1 hour later. Scale bar = 100 μm (*n* = 4). **J-M.** Quantification of MPO-positive and Cit H3-positive areas in the kidney (n = 4, ****p < 0.0001).

**Supplement 2.**
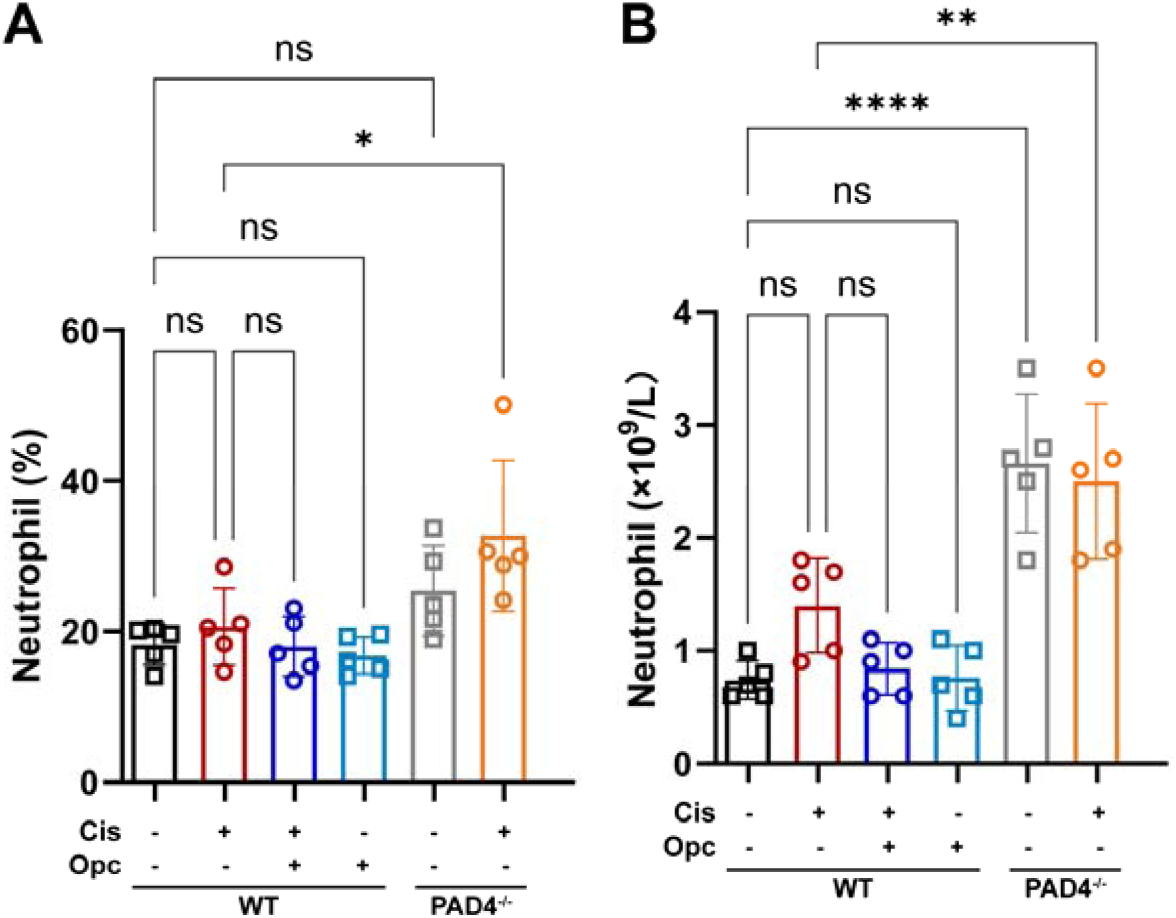
Number of neutrophils in the blood of mice from different groups. **A.** Quantitative analysis of the percentage of peripheral blood neutrophils in different groups of mice. **B.** Quantitative analysis of peripheral blood neutrophil counts in different groups of mice.

**Supplement 3.**
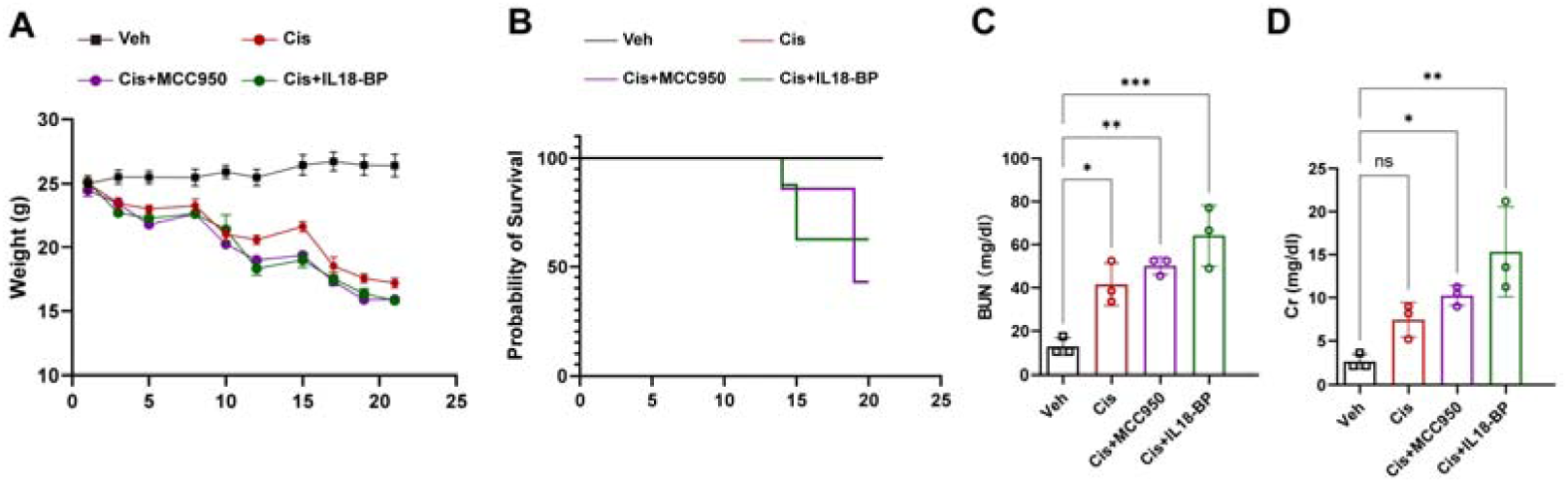
Status of RLDC modeling mice after anti-inflammatory treatment (Inhibition of corresponding inflammatory factors was achieved using the NLRP3 inhibitor MCC950 and the IL-18 inhibitor IL-18BP, followed by RLDC modelling for 21 days). **A.** The body weights of the mice during the three weeks of treatment (n = 10). **B.** Survival rate of mice during the three-week modeling period. **C, D.** Serum creatinine and blood urea nitrogen (n = 3).

**Supplement 4.**
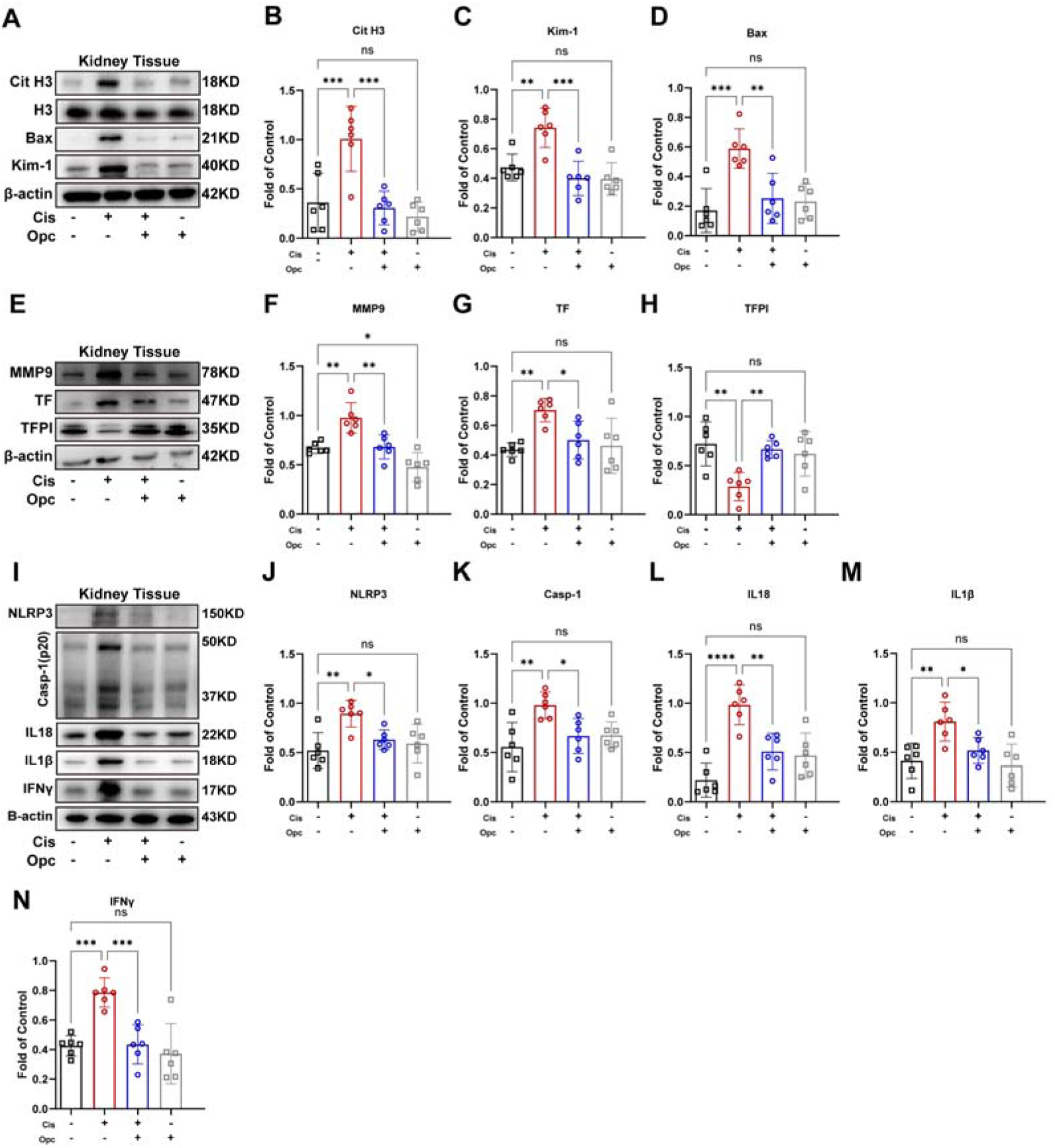
OPCs mitigate the formation of renal thrombosis and fibrosis caused by RLDC by inhibiting NETs. **A-N**. Immunoblot analysis of Cit H3, BAX, KIM-1, MMP9, TF, TFPI, NLRP3, Casp-1, IL18, IL1β and IFNγ in kidney tissues. For quantification, the protein was analyzed through densitometry and then normalized to β-actin (n = 6).

**Supplement 5.**
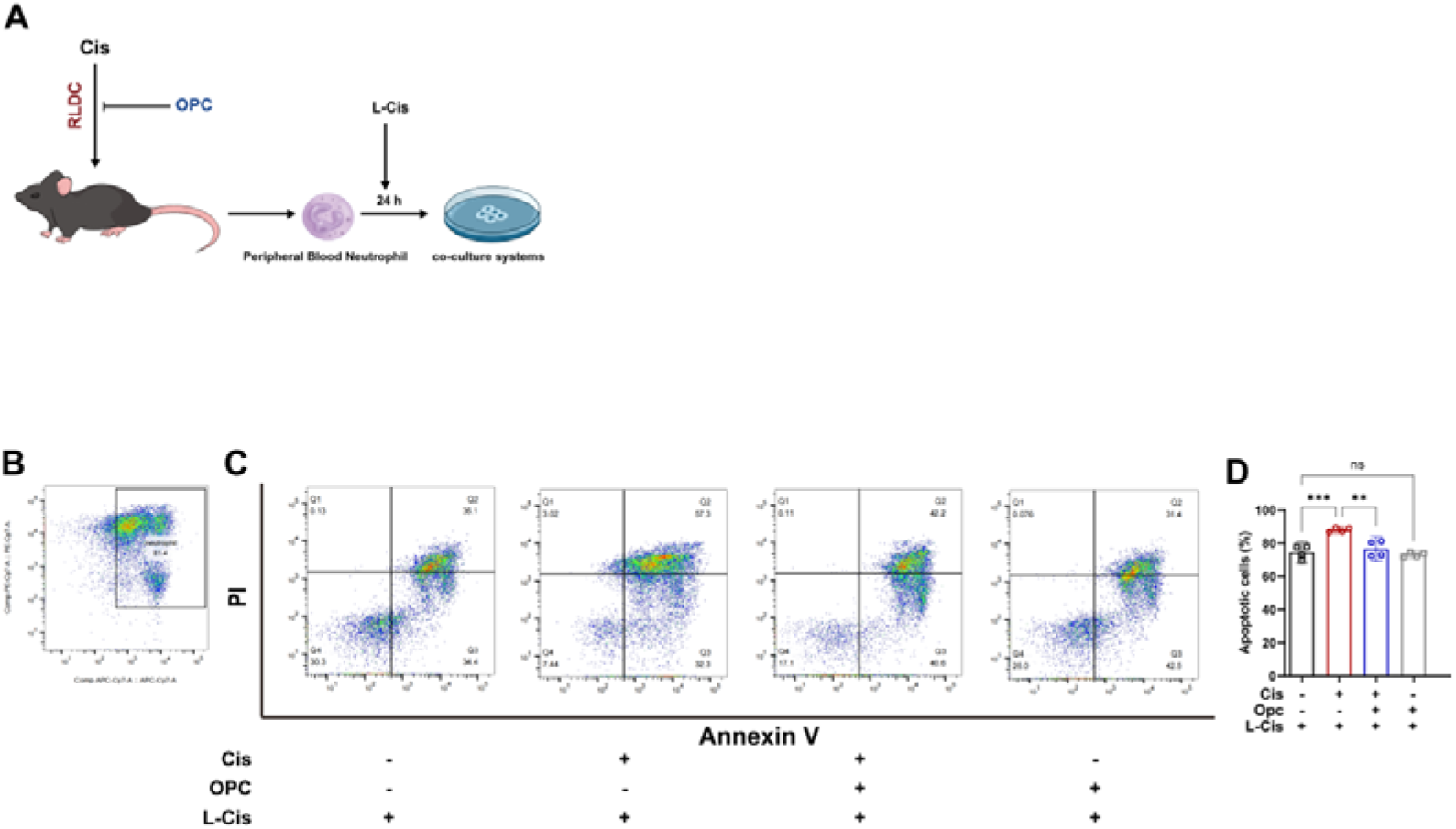
RLDC causes peripheral blood neutrophils in a hyperactive state while OPCs inhibit it. **A.** Illustrated explanation of peripheral blood neutrophils from different groups of mice, which aretreated with low-dose cisplatin. **B-D.** Apoptotic status of peripheral blood neutrophils 24 hours after low-dose cisplatin treatment (n = 4).

**Supplement 6.**
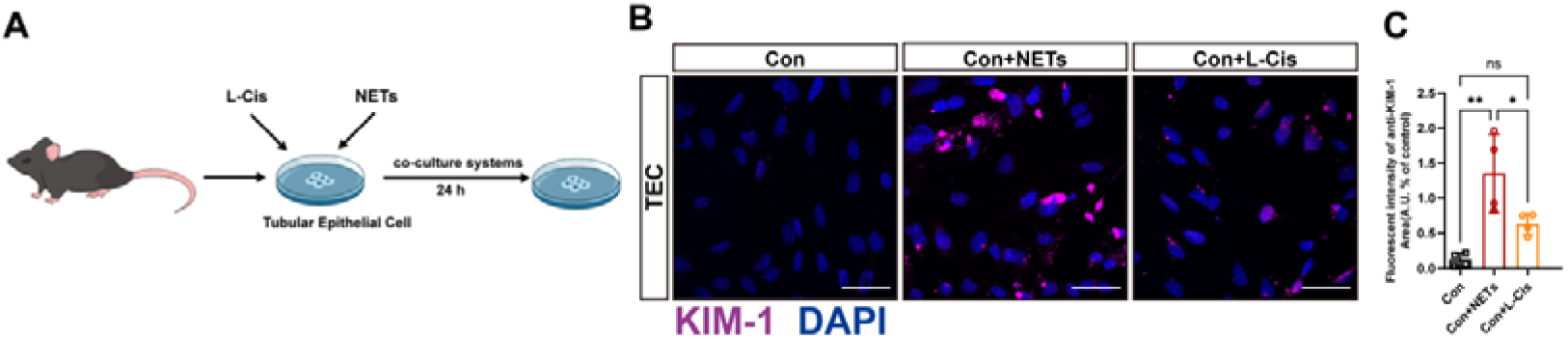
NETs are more likely to damage tubule cells than low-dose cisplatin. **A.** Illustration showing that renal tubular cells are treated with NETs or low-dose cisplatin for 24 hours. **B.** Immunofluorescence staining of primary renal tubule cells from mice. Anti-KIM-1 (purple), and DAPI (blue) staining are used to assess damage to tubule epithelial cells. **C.** Quantification of KIM-1-positive areas in tubule epithelial cells (n = 4, ****p < 0.0001).

**Supplement 7.**
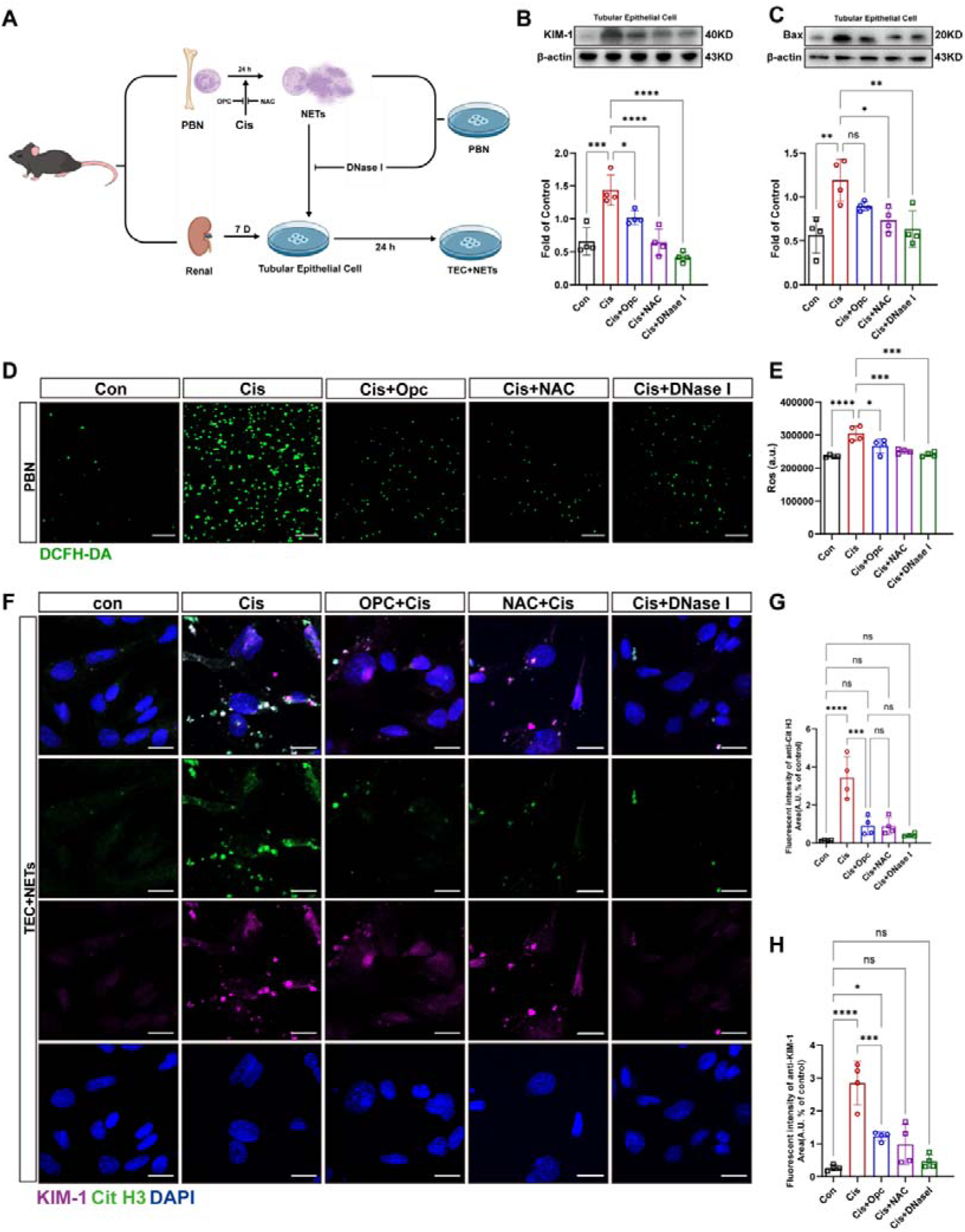
OPCs inhibit NETs damage to renal tubular epithelial cells. **A.** The diagram shows that peripheral blood neutrophils from mice in different groups are treated with primary renal tubular epithelial cells for 24 hours, which are subjected to different treatments. **B, C.** Immunoblotting analysis of BAX and KIM-1 in renal tubular epithelial cells from different treatment groups. For quantification, the protein is analyzed through densitometry and then normalized to β-actin (n = 4). **D.** Analysis of ROS in neutrophils from different treatment groups using DCFH-DA staining. **E.** Quantification of DCFH-DA staining of neutrophils by luminescence zymography (n = 4). F. Immunofluorescence staining of primary renal tubule cells from mice. Anti-Cit H3 (green), anti-KIM-1 (purple), and DAPI (blue) staining are used to assess damage to tubule epithelial cells. **G, H.** Quantification of KIM-1-positive areas in tubule epithelial cells (n = 4, ****

